# Tiny Earth CURE improves student persistence in science

**DOI:** 10.1101/2023.06.21.543782

**Authors:** Sarah Miller, Cristian Cervantes Aldana, Wenyi Du, Hyewon Lee, Natalia Maldonado, Perla Sandoval, Janice Vong, Gerald Young, Jo Handelsman, Nichole A. Broderick, Paul R. Hernandez, Mica Estrada

## Abstract

Course-based undergraduate research experiences (CUREs) enhance student retention in STEM, particularly among students who belong to historically excluded communities. Yet the mechanisms by which CUREs contribute to student integration and persistence are poorly understood. Utilizing the Tripartite Integration Model of Social Influence (TIMSI), this longitudinal study examines how Tiny Earth, an antibiotic-discovery CURE, impacts students’ scientific self-efficacy, scientific identity, endorsement of scientific community values, and intentions to persist in science. The study also explores how gains in TIMSI factors vary as a function of student demographics and course characteristics. Results of pre-and post-course measurements show that scientific self-efficacy and identity increased among students in Tiny Earth, and some student demographics and course characteristics moderated these gains. Gains in scientific self-efficacy, identity, and values correlated with gains in persistence intentions, whereas student demographics and course characteristics did not. Results of this study show that the Tiny Earth CURE enhanced students’ integration into the scientific community, which was linked to intentions of students of both historically underrepresented and majority groups to persist in STEM. We discuss how courses that provide opportunity to learn science skills in the context of a CURE can contribute toward enlarging and diversifying the STEM workforce.

## INTRODUCTION

To achieve a more vibrant and talented STEM workforce, it is imperative to recruit and retain diverse students in STEM. The United States currently wastes significant scientific talent by discouraging members of marginalized communities from pursuing STEM careers. Moreover, diversity within groups has been shown to increase intellectual rigor and creativity and quality of decisions, illustrating some of the benefits of diversifying the STEM workforce (Guimerà et al., 2005; McLeod et al., 1996; C. Nemeth et al., 2001; C. J. Nemeth & Kwan, 1987; Paulus et al., 2016; Sommers, 2006). Yet talented college students continue to leave STEM majors at disheartening rates, and those who are members of historically excluded communities (HECs) leave at higher rates than majority students (Thiry et al., 2019). Most of those who leave have the interest, confidence, and aptitude to be successful in STEM, but early classroom experiences dampen their interest and can be actively exclusive (Estrada et al., 2019; Thiry et al., 2019). Their departures after STEM introductory courses represent a major talent drain from the system.

Although students’ reasons for leaving have been studied extensively (Estrada et al., 2011; Estrada, Eroy-Reveles, et al., 2018; Rosenzweig et al., 2021), a stubborn gap persists in understanding the mechanisms by which certain interventions reduce attrition among students from HECs in STEM. Considerable research has shown that not all students integrate into their scientific community at the same rate and that three social influence mechanisms predict persistence in STEM (Estrada et al., 2011; Estrada, Hernandez, et al., 2018). Three factors contribute to integration: scientific self-efficacy, scientific identity, and internalization of scientific values. Scientific **self-efficacy** indicates that a student feels capable of performing the actions needed to be successful in a STEM course, major, or career. **Scientific identity** indicates that a student perceives science as part of their identity and feel they belong to a scientific community. Students internalize **scientific values** when they authentically agree with the values of the scientific community, such as building new knowledge to solve global challenges, the thrill of discovery, and the importance of discourse. According to the Tripartite Integration Model of Social Influence (TIMSI), each of these three factors—independently and collectively—contribute to social integration into STEM communities. **Social integration,** or **persistence,** is defined as the **intent** to pursue further academic or career goals in STEM, or as **behaviors**, such as participating in college research experiences and submitting applications for graduate school (Estrada et al., 2011).

The encouraging news is that education research has identified several practices that improve STEM performance and retention for students from HECs. One of the most effective ways to improve STEM outcomes and diversify the STEM student body is to focus on improving the situational factors connected to scientific self-efficacy, identity, and values. Course-based undergraduate research experiences (CUREs) are interventions that address many of these situational factors because they engage students with active learning and relevant content to increase efficacy, provide students a sense of belonging in STEM, and introduce students to the value of participating in science by engaging in meaningful data collection (National Academies of Sciences, Engineering, and Medicine, 2017).

### The Pedagogy of CUREs

Active learning has long been known to enhance the performance and retention of all students and is especially beneficial to students from HECs (Freeman et al., 2014; Haak et al., 2011; Hake, 1998; Hood et al., 2020; Schwartz et al., 2011; Theobald et al., 2020). According to Freeman et al. (2014), “Active learning engages students in the process of learning through activities and/or discussion in class, as opposed to passively listening to an expert. It emphasizes higher-order thinking and often involves group work.” Recent research has shined a light on the feasibility and importance of incorporating active-learning practices that have long-lasting effects on students and, in particular, those practices that reduce the inequities that result from students feeling disengaged and excluded from STEM (Freeman et al., 2014; Theobald et al., 2020).

**Course-based undergraduate research experiences** package the best of active learning and scientific objectives together, with a phenomenal track record of improving retention of diverse students in STEM. Although each CURE is unique in scope and format, CUREs commonly center on iterative scientific research that addresses a relevant problem. CUREs give students ownership of their experiments, the opportunity to disseminate their results, and the chance to make a new discovery. Evidence suggests that CUREs improve outcomes for all students (Rodenbusch et al., 2016) and can have especially positive effects on students from HECs (Evans et al., 2021; Hurtado et al., 2009; Olson et al., 2019; Shuster et al., 2019; Waddell et al., 2021). CUREs enable students to be scientists in a community of peers, thereby increasing their opportunity to identify as scientists, regardless of ethnicity or background. CUREs also make research experiences available to all students on an equal basis (Hanauer et al., 2022; Hurtado et al., 2009). In one study, students in CUREs gained in scientific self-efficacy, scientific identity, and career intent significantly more than students in a non-CURE control group (Newell & Ulrich, 2022).

CUREs scale better than research experiences in faculty research labs. The course-based nature of CUREs means that they can be available to all students, regardless of background or experience, whereas faculty research labs are small, exclusive, and typically limited to experienced students. At some undergraduate institutions, such as community colleges, faculty research programs are typically not an option, and CUREs may be the only way for students to participate in research. In some cases, CUREs lead to more positive outcomes than research in faculty labs. For example, research in faculty labs can influence scientific identity and intent to pursue graduate study among women and Asian students less positively than members of other demographic groups (Aikens et al., 2017), whereas CUREs have positive effects on students of all demographic groups (Evans et al., 2021; Olson et al., 2019; Rodenbusch et al., 2016).

Several CURE programs, such as the Science Education Alliance – Phage Hunters Advancing Genomics and Evolutionary Science (SEA-PHAGES) program, the Genomics Education Partnership (GEP), and UCLA’s Undergraduate Research Consortium in Functional Genomics, set a standard for quality and impact, demonstrating that research courses can nurture creativity, reinforce diverse talents, diminish the stigma of failure, foster community, and teach principles of equity and inclusion in science (Evans et al., 2021; Hanauer et al., 2017; Jordan et al., 2014; Lopatto et al., 2008; Olson et al., 2019; Shaffer et al., 2014; Waddell et al., 2021). They also provide students an opportunity to take project ownership and develop identities as scientists in a community of peers (Hanauer et al., 2017). Studies show that taking a CURE early in college increases the likelihood that a student will remain in a STEM major until graduation and that they will complete college, regardless of major (Rodenbusch et al., 2016), suggesting that CUREs have a significant impact on students’ academic trajectories well beyond the research course itself.

Despite the promising outcomes of CUREs, there is a gap in understanding the mechanisms underlying these benefits. In this paper, we will apply the Tripartite Integration Model of Social Influence (Estrada et al., 2011) to understand whether and how gains in scientific self-efficacy, identity, and values orientation contribute to gains in social integration into the scientific community experienced by students in the Tiny Earth CURE at 25 U.S. colleges and universities.

### The Tiny Earth CURE

Tiny Earth is a new, international approach to antibiotic discovery. Tiny Earth aims to address a pressing global health crisis, antibiotic resistance, which is predicted to cause 50 million deaths per year by 2050 (World Health Organization, n.d.). People need a new approach to antibiotic discovery because no new structural classes of antibiotics have been discovered since the 1980s. The Tiny Earth CURE leverages an untapped natural resource – soil bacteria from diverse locations – and an untapped workforce – college students – to crowdsource antibiotic discovery. An estimated 14,000+ students per year in 30 countries enroll in a Tiny Earth CURE, each of which is taught by a trained instructor at their college or university. The students’ collective search forms a massive network focused on antibiotic discovery. Tiny Earth’s “studentsourcing” approach could provide an antidote to the abandonment of antibiotic discovery by the pharmaceutical industry, which instead focuses on synthetic derivatives of known antibiotics or treatments for chronic diseases, which are more profitable than infectious disease. Tiny Earth makes research accessible to students at any type of college, including minority-serving institutions, community colleges, 4-year colleges, and research universities (S. Hernandez et al., 2020; Hurley et al., 2021).

To take advantage of the many pedagogical and inclusive features of CUREs, Tiny Earth embodies the six hallmarks of a CURE: scientific research, iteration, ownership, relevance, discovery, and dissemination (Dolan & Weaver, 2021). Students choose their own soil, growth media, and research questions; engage in replicated, hypothesis-driven research; discover antibiotic-producing bacteria from soil; and communicate their research findings. The goal of Tiny Earth is to discover new antibiotics from soil bacteria and encourage diverse students to persist in STEM. **Box 1** shows recommended student learning objectives for the Tiny Earth course. In 2020, modifications were made for teaching remotely (González-Orta et al., 2022) and with content based on principles of antiracism, justice, equity, diversity, and inclusion (AJEDI) (Miller et al., 2022).

A Tiny Earth instructor training program, based on the National Institute on Scientific Teaching model (Handelsman et al., 2004, 2007; Miller et al., 2008; Pfund et al., 2009), immerses participants in pedagogical principles of scientific teaching along with the experimental content of the Tiny Earth CURE. Since 2013, more than 700 instructors have been trained who teach an estimated 14,000+ students per year. **Box 2** presents a typical training schedule for instructors.

### Research Theory

The guiding theory for this research is the Tripartite Integration Model of Social Influence (TIMSI) (Estrada, Eroy-Reveles, et al., 2018; Estrada et al., 2011). The TIMSI is a model of social influence that describes how a person’s orientation toward a social system can predict the conditions under which they would conform to the demands of the influencing agents, such as instructors, mentors, peers, or the design of learning contexts. TIMSI describes three orientations that influence persistence outcomes: efficacy, identity, and values. Previous TIMSI research has examined the impact of science training programs and mentorship on integration and long-term persistence for STEM students who are from HECs and historically included communities (HICs) (Estrada, Eroy-Reveles, et al., 2018; Estrada et al., 2011; P. R. Hernandez et al., 2018, 2020). This study tests TIMSI-based research questions in a national sample of undergraduates in the Tiny Earth CURE at 25 colleges (*k* = 31 instances of the course). The current study extends prior research by longitudinally tracking TIMSI-identified factors from the beginning through the end of a one-semester Tiny Earth course, with a large enough sample of courses to examine the impact of course characteristics in addition to student individual differences, such as gender, race and ethnicity, and year in school.

### Research Questions and Hypotheses

While participating in a Tiny Earth CURE:

1. What, if any, gains do students experience in the TIMSI mechanisms of scientific self-efficacy, identity, or values? We hypothesized that the Tiny Earth CURE positively affects students’ scientific self-efficacy, identity, and values orientation (H1).
2. What, if any, student demographics or course characteristics influence gains in the TIMSI mechanisms? We hypothesized that Tiny Earth positively affects TIMSI mechanisms across all student demographics (H2a). Furthermore, we hypothesized that Tiny Earth positively affects TIMSI mechanisms across all course characteristics (H2b).
3. What, if any, gains do students experience in their intent to persist in science careers? We hypothesized that Tiny Earth positively affects students’ intent to persist in science careers (H3).
4. In what ways do student characteristics, course characteristics, and/or gains in TIMSI mechanisms impact the relationship between participation in Tiny Earth and gains in student persistence intentions?

Answers to these research questions will advance knowledge regarding whether students are more likely to persist when they participate in a Tiny Earth CURE as well as “why?” and “for whom?”

## METHODS

### Participants

The current study used data from a large, national longitudinal study of undergraduate students taking a course that incorporated the Tiny Earth curriculum. The data were collected from 1,419 undergraduate students grouped into 33 biological science college classrooms across the United States. Two classes with 121 participants were excluded from the current study as their instructors did not meet the criteria of significantly incorporating Tiny Earth elements into their teaching. Participants were omitted from the analytic sample if they did not respond to the end-of-semester survey (*n* = 502), did not respond to survey questions concerning the relevant variables (*n* = 49), were not an undergraduate while enrolled in the course (*n* = 46), or provided inconsistent responses to survey questions (e.g., selecting all possible racial and/or gender categories; *n* = 3). As a result, the final analytical sample included 698 students grouped within 31 Tiny Earth courses at 25 colleges and universities. The majority self-identified as female (52%). The largest racial and ethnic groups were of white, non-Hispanic (48%), Latine (18%), Asian (13%), or multi-racial (13%) descent. Nearly half were first-generation college students (47%), and most were in their sophomore (35%), junior (24%), or senior (27%) year of college (see **Table 1** for complete demographic data). Most of the courses were low-enrollment (i.e., *n_j_* < 20; 45%) or medium-enrollment (i.e., 20 ≤ *n_j_* < 100; 42%). Courses were evenly split between lower-or upper-division. Most (65%) were taught in-person, and the rest were hybrid; none were taught fully online during the semester of the study. The majority (68%) included a combination of lecture and laboratory components, and the rest were classified as lab-only courses. Most instances of the Tiny Earth course integrated the curriculum into a microbiology course (61%), whereas several others integrated it into a biology course (19%) or taught it as a standalone course (16%; see **Table 2**).

**Table 1.**
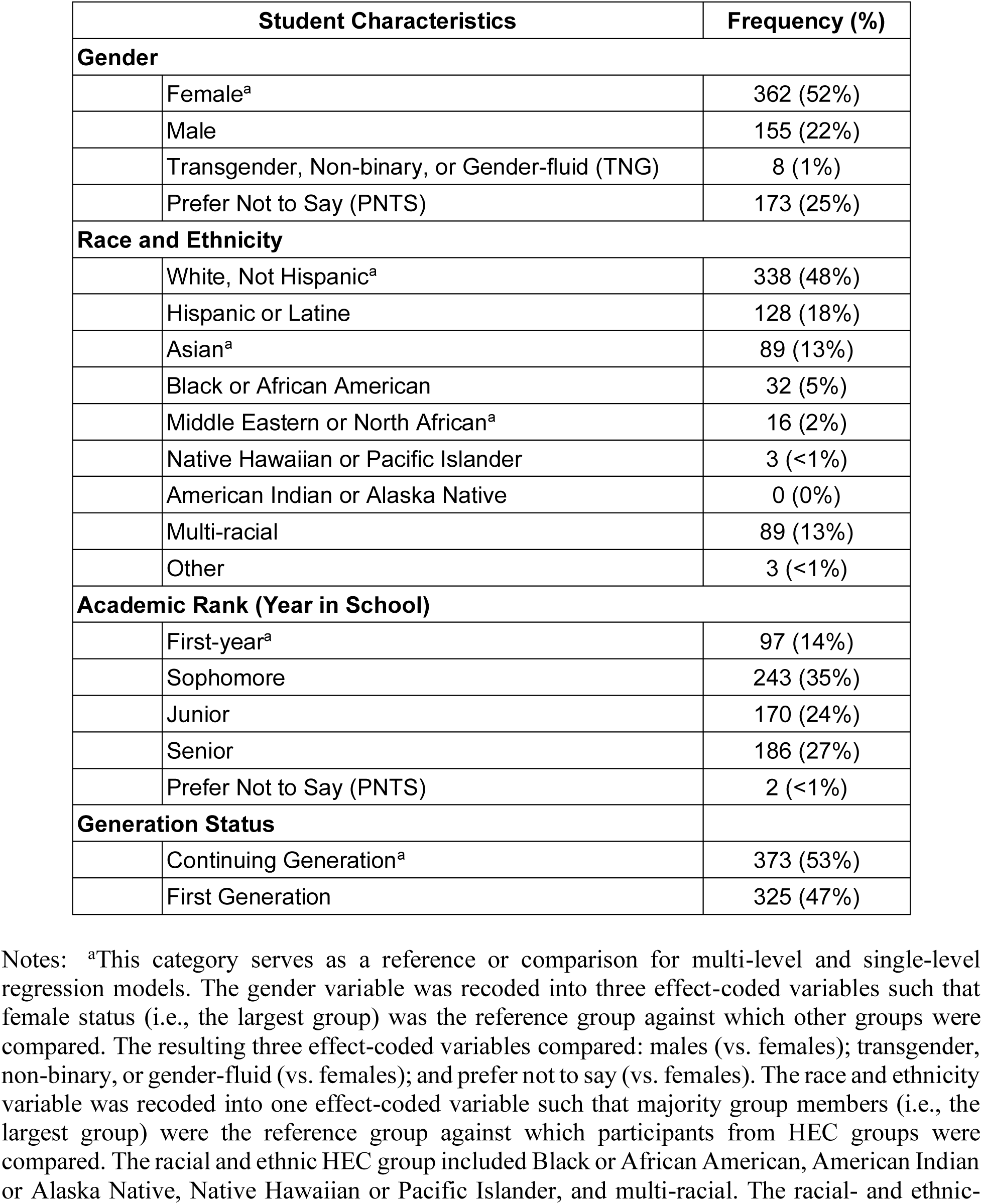
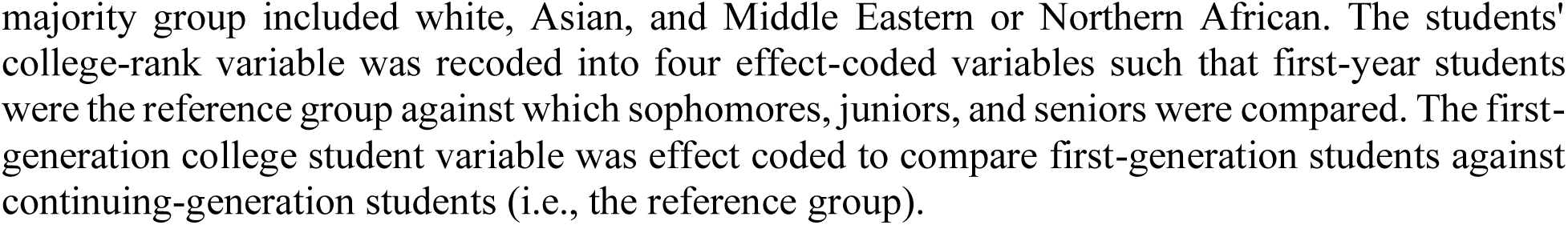
Tiny Earth student demographics (*N*=698, *J*=31, *n_j_*=22.52). Demographic information about Tiny Earth students who responded to both pre-and post-surveys includes gender identity, racial and ethnic identity, academic rank (year in school), and continuing-or first-generation status (*N*=698).

| Student Characteristics |  | Frequency (%) |
| --- | --- | --- |
| **Gender** |  |  |
|  | Female<sup>a</sup> | 362 (52%) |
|  | Male | 155 (22%) |
|  | Transgender, Non-binary, or Gender-fluid (TNG) | 8 (1%) |
|  | Prefer Not to Say (PNTS) | 173 (25%) |
| **Race and Ethnicity** |  |  |
|  | White, Not Hispanic<sup>a</sup> | 338 (48%) |
|  | Hispanic or Latine | 128 (18%) |
|  | Asian<sup>a</sup> | 89 (13%) |
|  | Black or African American | 32 (5%) |
|  | Middle Eastern or North African<sup>a</sup> | 16 (2%) |
|  | Native Hawaiian or Pacific Islander | 3 (<1%) |
|  | American Indian or Alaska Native | 0 (0%) |
|  | Multi-racial | 89 (13%) |
|  | Other | 3 (<1%) |
| **Academic Rank (Year in School)** |  |  |
|  | First-year<sup>a</sup> | 97 (14%) |
|  | Sophomore | 243 (35%) |
|  | Junior | 170 (24%) |
|  | Senior | 186 (27%) |
|  | Prefer Not to Say (PNTS) | 2 (<1%) |
| **Generation Status** |  |  |
|  | Continuing Generation<sup>a</sup> | 373 (53%) |
|  | First Generation | 325 (47%) |

Notes: <sup>a</sup>This category serves as a reference or comparison for multi-level and single-level regression models. The gender variable was recoded into three effect-coded variables such that female status (i.e., the largest group) was the reference group against which other groups were compared. The resulting three effect-coded variables compared: males (vs. females); transgender, non-binary, or gender-fluid (vs. females); and prefer not to say (vs. females). The race and ethnicity variable was recoded into one effect-coded variable such that majority group members (i.e., the largest group) were the reference group against which participants from HEC groups were compared. The racial and ethnic HEC group included Black or African American, American Indian or Alaska Native, Native Hawaiian or Pacific Islander, and multi-racial. The racial- and ethnic-
majority group included white, Asian, and Middle Eastern or Northern African. The students' college-rank variable was recoded into four effect-coded variables such that first-year students were the reference group against which sophomores, juniors, and seniors were compared. The first-generation college student variable was effect coded to compare first-generation students against continuing-generation students (i.e., the reference group).

**Table 2.**
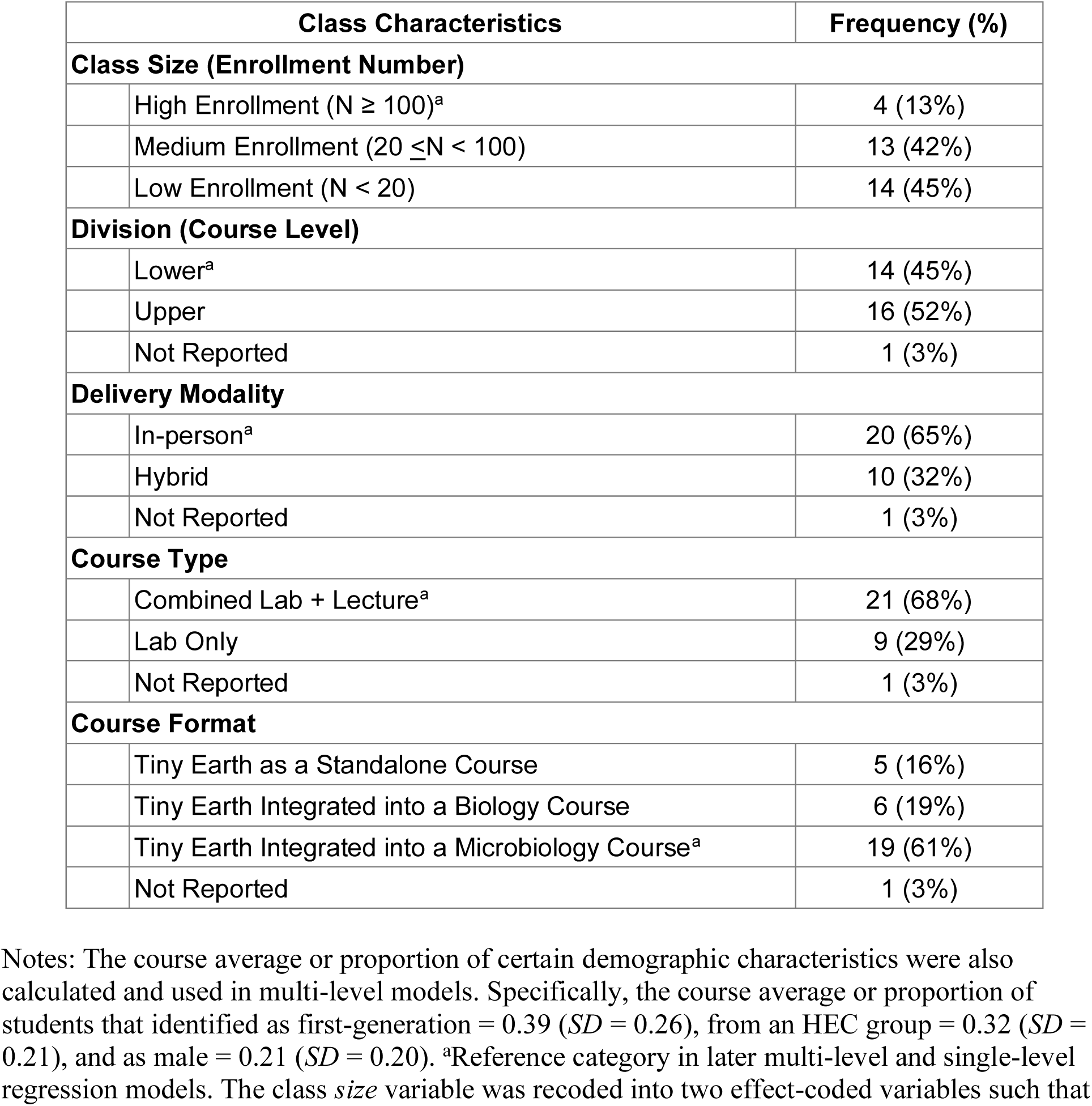
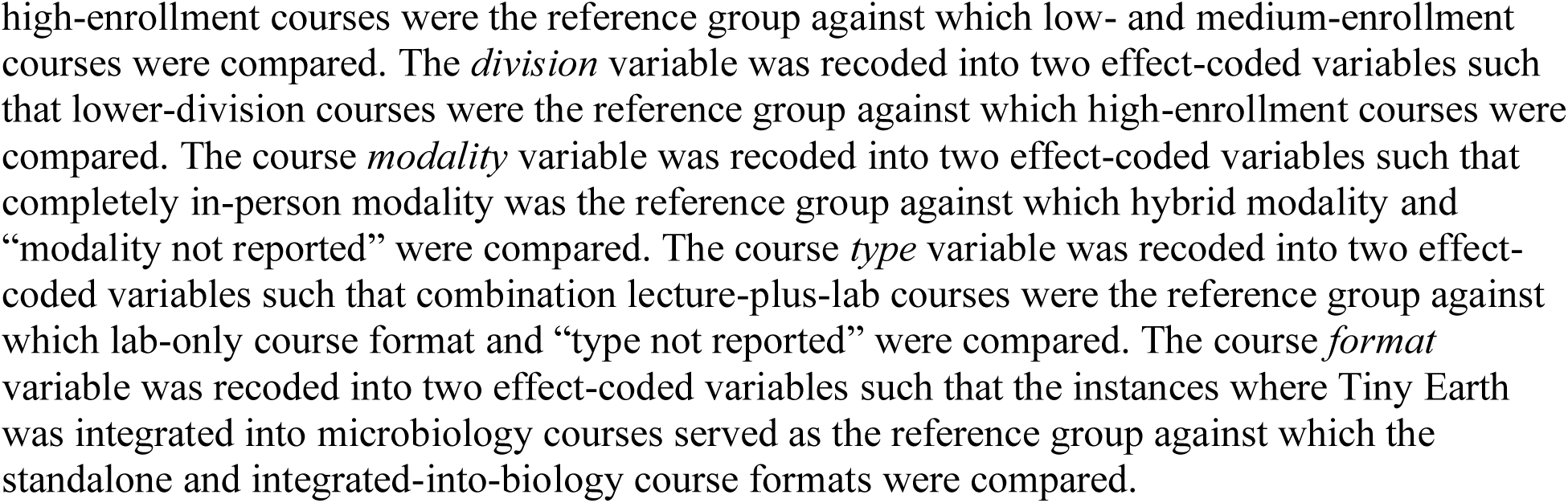
Tiny Earth course characteristics (*J*=31). Course characteristics for the Tiny Earth courses in the study included class size, division, delivery modality, type, and format. Class sizes were categorized into low enrollment (*N* < 20), medium enrollment (20 ≤ *N* < 100), and high enrollment (*N* ≥ 100). Division categories described the course level, indicating whether the course was intended as lower-division course for first-or second-year students, or an upper-division course for junior or senior students. Delivery modality categories described the extent to which the course was fully in person, fully remote, or a combination (‘hybrid’). Course type categories differentiated which courses were taught as a laboratory-only configuration or part of a lecture-plus-laboratory combination. The format category designated whether Tiny Earth was taught as a standalone course (e.g., a first-year research seminar or independent study) or integrated into another microbiology or biology course (e.g., as the part of an introductory biology curriculum) (*J* = 31).

| Class Characteristics |  | Frequency (%) |
| --- | --- | --- |
| **Class Size (Enrollment Number)** |  |  |
| | High Enrollment ($N \geq 100$)<sup>a</sup> | 4 (13%) |
| | Medium Enrollment ($20 \leq N < 100$) | 13 (42%) |
| | Low Enrollment ($N < 20$) | 14 (45%) |
| **Division (Course Level)** |  |  |
|  | Lower<sup>a</sup> | 14 (45%) |
|  | Upper | 16 (52%) |
|  | Not Reported | 1 (3%) |
| **Delivery Modality** |  |  |
|  | In-person<sup>a</sup> | 20 (65%) |
|  | Hybrid | 10 (32%) |
|  | Not Reported | 1 (3%) |
| **Course Type** |  |  |
|  | Combined Lab + Lecture<sup>a</sup> | 21 (68%) |
|  | Lab Only | 9 (29%) |
|  | Not Reported | 1 (3%) |
| **Course Format** |  |  |
|  | Tiny Earth as a Standalone Course | 5 (16%) |
|  | Tiny Earth Integrated into a Biology Course | 6 (19%) |
|  | Tiny Earth Integrated into a Microbiology Course<sup>a</sup> | 19 (61%) |
|  | Not Reported | 1 (3%) |

Notes: The course average or proportion of certain demographic characteristics were also calculated and used in multi-level models. Specifically, the course average or proportion of students that identified as first-generation = 0.39 ( $SD = 0.26$ ), from an HEC group = 0.32 ( $SD = 0.21$ ), and as male = 0.21 ( $SD = 0.20$ ). <sup>a</sup>Reference category in later multi-level and single-level regression models. The class *size* variable was recoded into two effect-coded variables such that
high-enrollment courses were the reference group against which low- and medium-enrollment courses were compared. The *division* variable was recoded into two effect-coded variables such that lower-division courses were the reference group against which high-enrollment courses were compared. The course *modality* variable was recoded into two effect-coded variables such that completely in-person modality was the reference group against which hybrid modality and “modality not reported” were compared. The course *type* variable was recoded into two effect-coded variables such that combination lecture-plus-lab courses were the reference group against which lab-only course format and “type not reported” were compared. The course *format* variable was recoded into two effect-coded variables such that the instances where Tiny Earth was integrated into microbiology courses served as the reference group against which the standalone and integrated-into-biology course formats were compared.

### Procedure

College biology and microbiology instructors were recruited from a national network of instructors to apply to attend a week-long training program about teaching the Tiny Earth CURE. Instructors who intended to implement Tiny Earth as a course at their college or university were selected from the pool of applicants. Student recruitment to the study involved providing the instructor with a Google slide containing a brief study description, along with an anonymous Qualtrics survey link. Each instructor was asked to distribute our study invitation to their Tiny Earth class of students (e.g., via email or learning management system).

The students completed a pre-course survey at the beginning of one semester (T1) and a post-course survey at the end of that same semester (T2). As an incentive, participants received a $10 gift card for completing each survey. Student participants were recruited into the study in four cohorts during the Fall of 2020, Spring of 2021, Fall of 2021, and Spring of 2022. The study was approved by the UCSF Institutional Review Board (IRB# 19-28867) and in consultation with all institutions in which data were collected. Surveys are included in **Supplemental Materials**.

### Measures

#### Scientific Self-efficacy

We used a three-item scale adapted from a previous study (Estrada et al., 2011) to assess the students’ confidence in their ability to function as a scientist in a variety of tasks (e.g., “Create explanations for the results of the study”). Student participants rated each of the statements with a five-point Likert scale from 1 (not at all confident) to 5 (absolutely confident). Responses were averaged to derive composite scores to indicate their overall scientific self-efficacy at each time point (*α_T1_* = 0.84 and *α_T2_* = 0.88). A gain score was calculated as the difference between the participants’ pre-and post-course survey responses (i.e., positive scores indicate higher self-efficacy post-course than pre-course).

#### Identification as a Scientist

We utilized a four-item scientific-identity scale adapted from a previous study (Estrada et al., 2011) to assess the degree to which the participants identified as scientists. Participants were asked to assess on a Likert scale of 1 (strongly disagree) to 5 (strongly agree) their agreement with a series of statements (e.g., “Being a scientist is an important reflection of who I am”). Responses were averaged to derive composite scores to indicate their overall scientific identity at each time point (*α_T1_* = 0.88 and *α_T2_* = 0.88). A gain score was calculated as the difference between the participants’ pre-and post-survey.

#### Internalization of Scientific Community Values

We used a four-item scale from a previous study (Estrada et al., 2011) to measure the extent to which participants endorsed values of the scientific community, such as discovery and experimentation. Participants rated the degree to which statements applied to them (e.g., “A person who thinks it is valuable to conduct research that builds the world’s scientific knowledge”) on a scale from 1 (not at all like me) to 6 (very much like me). Responses were averaged to derive composite scores to indicate their overall scientific values at each time point (*α_T1_* = 0.87 and *α_T2_* = 0.88). A gain score was calculated as the difference between the participants’ pre-and post-course survey.

#### Scientific Career Persistence Intentions

A seven-item scale adapted from previous research (Woodcock et al., 2012) was used to assess participants’ intention to persist in a scientific career. Participants rated their intention to persist (e.g., “To obtain a biomedical science-related undergraduate degree”) on a scale from 0 (definitely will not) to 10 (definitely will). Responses were averaged to derive a composite score (*α_T1_* = 0.84 and *α_T2_* = 0.86). A gain score was calculated as the difference between the participants’ pre-and post-course survey.

#### Student Characteristics

The participants were asked to provide their demographic information for four categories: self-identified gender, race and ethnicity, current year in school, and first-generation college-student status **(Table 1**). All categorical variables were recoded for the analysis to facilitate comparisons with a reference group (e.g., each gender group was compared to the female reference group) and to facilitate interpretation of the regression intercept as the average across groups and classes. To achieve these analysis goals, we recoded each categorical variable into a set of effect-coding group comparison variables, such that the comparison group was coded 0.5, and all others were coded as -0.5 (Yaremych et al., 2021).

For analysis, responses were grouped into categories. Gender analysis categories included four groups: women; men; students who identified as transgender, non-binary, or gender-fluid (TNG); and those who preferred not to say their gender (PNTS). For race and ethnicity analysis, students were grouped into two categories: HEC and HIC. The HEC group included those who identified as Black or African American, Hispanic or Latine, American Indian or Alaska Native, Native Hawaiian or Pacific Islander, and multi-racial. More than half of the multi-racial group (*n = 33*) also included those who selected both white and Hispanic/Latine. The racial and ethnic HIC group included white (not Hispanic), Asian, and Middle Eastern or Northern African. Students were also grouped according to first-or continuing-generation status and by year in school (first-year, sophomore, junior, or senior).

#### Course Characteristics

Participants were nested within 31 Tiny Earth classes. Information on five class-level variables (size, level, modality, type, and format) was collected directly from instructors for the semester in which their Tiny Earth course was included in the study **(Table 2)**. As with student characteristics, course-characteristic categorical variables were recoded into a set of effect-coding group comparison variables so that the comparison group was coded 0.5 and all others were coded -0.5.

Courses were classified based on *size*, with those containing fewer than 20 students considered low enrollment, those with 20 or more but less than 100 considered medium enrollment, and those with 100 or more considered high enrollment. The *division* variable distinguished lower-division courses, which were intended for first-and second-year students, from upper-division courses, which were intended for junior and senior students. The delivery *modality* variable distinguished between courses that were completely in-person, those that used a hybrid modality (i.e., partially in-person and partially online/remote), and those that were completely remote or online. No courses reported being completely online/remote only; therefore this category was not considered or reported further. The course *type* variable distinguished courses that were taught in a laboratory-only format from those that combined lectures with a laboratory component. Course *format* differentiated Tiny Earth courses taught as standalone courses (e.g., as a first-year seminar or independent research credit) from those integrated into another biology or microbiology course (e.g., the lab component of an introductory biology course).

We also derived *course-level demographic characteristics* for each class by calculating the proportion of students from focal demographic characteristics (i.e., proportion first-generation, proportion from an HEC group, and proportion female in each class). The proportion of transgender, non-binary, and gender-fluid participants in each class was not derived due to the extremely small sample size and preponderance of classes with 0%. Further, the proportion of ‘prefer not to say’ their gender in each class was not derived because this gender category was used as a control variable and does not have a directly interpretable meaning. Interestingly, 25% of students actively opted not to provide their gender identity. Similarly, year in school was treated as a control variable, and thus we did not aggregate this variable in order to distinguish within-and between-class effects.

#### Statistical Assumption Tests

All analyses were conducted in Stata v.17 (StataCorp, 2021)). We conducted preliminary analysis of statistical assumption checks and missing data analysis to (1) understand whether our statistical models would produce accurate and unbiased statistical results, (2) test longitudinal measurement properties of outcomes to determine whether the outcomes exhibited consistent measurement properties over time, and (3) test the cross-group measurement invariance of all outcomes at each time point as a function of gender and race/ethnicity to determine whether the outcomes exhibited consistent measurement properties across demographic groups. Overall, we concluded that these tests showed robust evidence that statistical assumptions were tenable and of longitudinal measurement invariance for each construct over time. Importantly, the assumption tests also showed that whereas gains scores in scientific self-efficacy, identity, and values exhibited small intraclass correlations (ICC; i.e., clustering effect that can be handled through multilevel modeling), gain scores in science persistence intentions exhibited an ICC so close to zero it caused convergence problems. Based on this information, we tested our research questions and hypotheses about scientific efficacy, identity, and values using multilevel regression models and tested our research questions about science persistence intentions using multiple regression models. The analyses and findings are fully described in **Supplemental Materials and Tables**.

#### Analytic Approach to Testing Hypotheses

To formally test research question and hypothesis 1 (*quantifying gains in TIMSI mechanisms*), we fitted a series of unconditional multilevel models (i.e., no predictors) to estimate the overall average gains in efficacy, identity, and values across classes.

To test research question 2 and hypotheses 2a and 2b (*testing student and course moderators of gains, respectively*), we fitted a series of multilevel models predicting student gain scores in efficacy, identity, and values from student demographics and course characteristics (Rights & Sterba, 2019). The focal within-classroom student demographic characteristics (i.e., female vs. male status, HEC vs. majority status, and first-vs. continuing-generation status) were group-mean centered within classes, as well as being aggregated to the classroom level (e.g., proportion of HEC students per class) and grand mean (Rights & Sterba, 2019; Yaremych et al., 2021). The advantage of our approach to coding, aggregating, and centering is that it allowed us to disentangle the within-classroom impact of demographic characteristics versus between-classroom impact of these characteristics. Further, this allowed us to interpret the effects of student demographic characteristics as the average group-mean difference in gains scores within classes (e.g., mean difference in scientific identity gains for women versus men in their classes, on average across classes). Second, it enabled us to interpret the effects of aggregated classroom-level demographic characteristics as the mean difference in gains scores for classes with 0% versus 100% of the demographic characteristic (e.g., mean difference in scientific identity for classes with no women versus all women). All other within-classroom student characteristics were treated as control variables and uncentered for analyses. For between-classroom course characteristics, the effect-coded categorical variables were uncentered and the continuous variables were grand-mean centered.

To test research question 3 and hypothesis 3 (*quantifying gains in science persistence intentions*), we fitted an unconditional multilevel model to estimate the overall average gains in science persistence intentions across classes.

To test research question 4 (*the extent to which student demographics, course characteristics, and the TIMSI mechanisms predict science persistence intentions*), we used a multiple regression model to predict gains in science persistence intentions from student demographics, course characteristics, and gains in student scientific self-efficacy, identity, and values. This model accounted for the lack of between-class variability in persistence intentions gain scores. For student and course characteristics, the categorical variables were effect coded, and for the TIMSI mechanisms, the continuous variables were centered. To understand the relationships, we first binned the gains in each of the TIMSI mechanisms into three groups: average gains, representing those who reported average gains in self-efficacy, identity, or values; low gains, representing the lowest (-1 SD) gains in the TIMSI mechanisms; or high gains (+1 SD), representing the highest gains in the mechanisms. These three bins were then mapped onto gains in persistence intentions.

## RESULTS

### Results of Testing Our Research Questions and Hypotheses

#### Hypothesis 1

*The Tiny Earth CURE positively affects students’ scientific self-efficacy, identity, and values orientation*. Overall, the starting course average levels of scientific efficacy, identity, and values were moderately high: Course averages ranged between 60-80% of the maximum possible score. After the course, multilevel regression models indicated *scientific self-efficacy* and *identity* showed significant average gain score across classes (consistent with H1), whereas average gain score in *scientific values* were negligible across classes (inconsistent with H1; **Table 3)**.

**Table 3.**
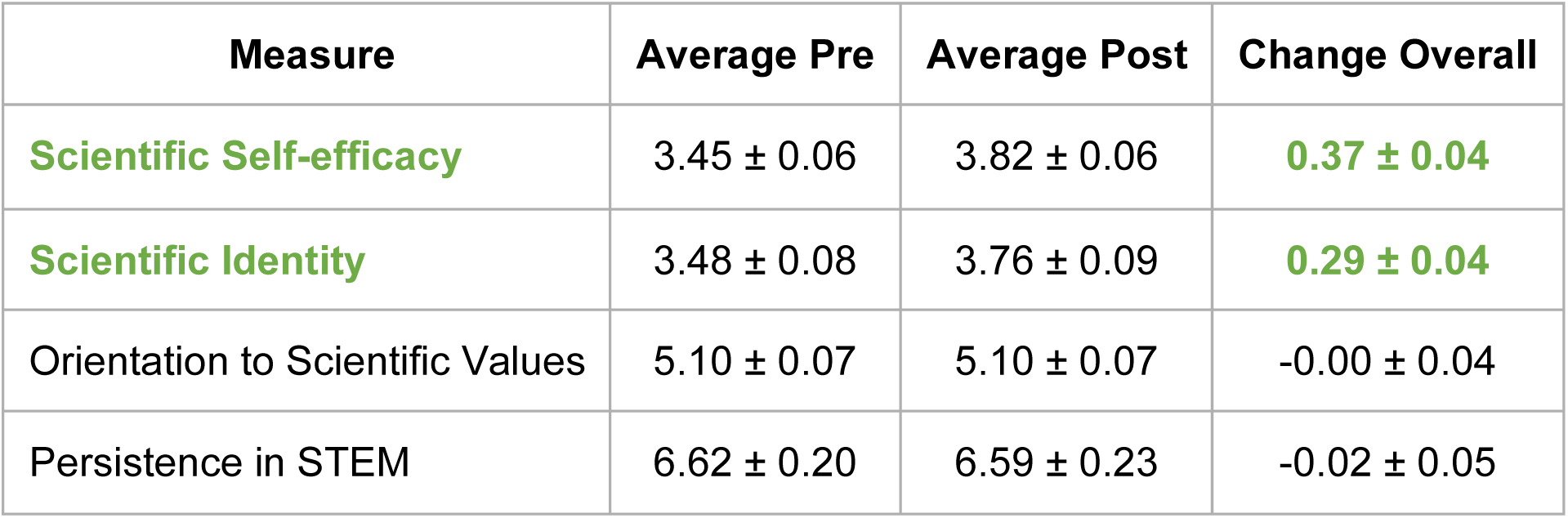
Overall impact of the Tiny Earth CURE on student’s scientific self-efficacy (E), identity (I), values (V) orientation, and persistence intentions (P). Using paired pre-and post-surveys, the impact of Tiny Earth was measured using the three orientations of the Tripartite Integration Model of Social Influence (TIMSI) that influence persistence in STEM: scientific self-efficacy; identity as a scientist; and orientation toward scientific community values, such as discovery and collaboration. Gains in E, I, V, and P were measured using a paired pre-and post-survey during the semester in which the Tiny Earth students’ course was included in the study. This figure shows the average, paired gains in E, I, V, and P for all students who responded to both surveys (*N = 698; p < 0.05*). Self-efficacy and identity were measured on scales from 1 to 5. Values were measured on a scale from 1 to 6. Persistence was measured on a scale from 1 to 10.

#### Hypothesis 2a

*Tiny Earth positively affects TIMSI mechanisms across all student demographics*. Next, we conducted a series of multilevel models predicting gain scores from student demographics and course characteristics. Consistent with our hypothesis, the analysis revealed all demographic groups exhibited significant gains in *scientific self-efficacy* and no meaningful group-mean differences between first-generation students and their continuing-generation classmates, women and their male classmates, or students from a HEC group and their majority classmates within classes (**Fig. 1A & Supplemental Tables**).

**Fig. 1.**
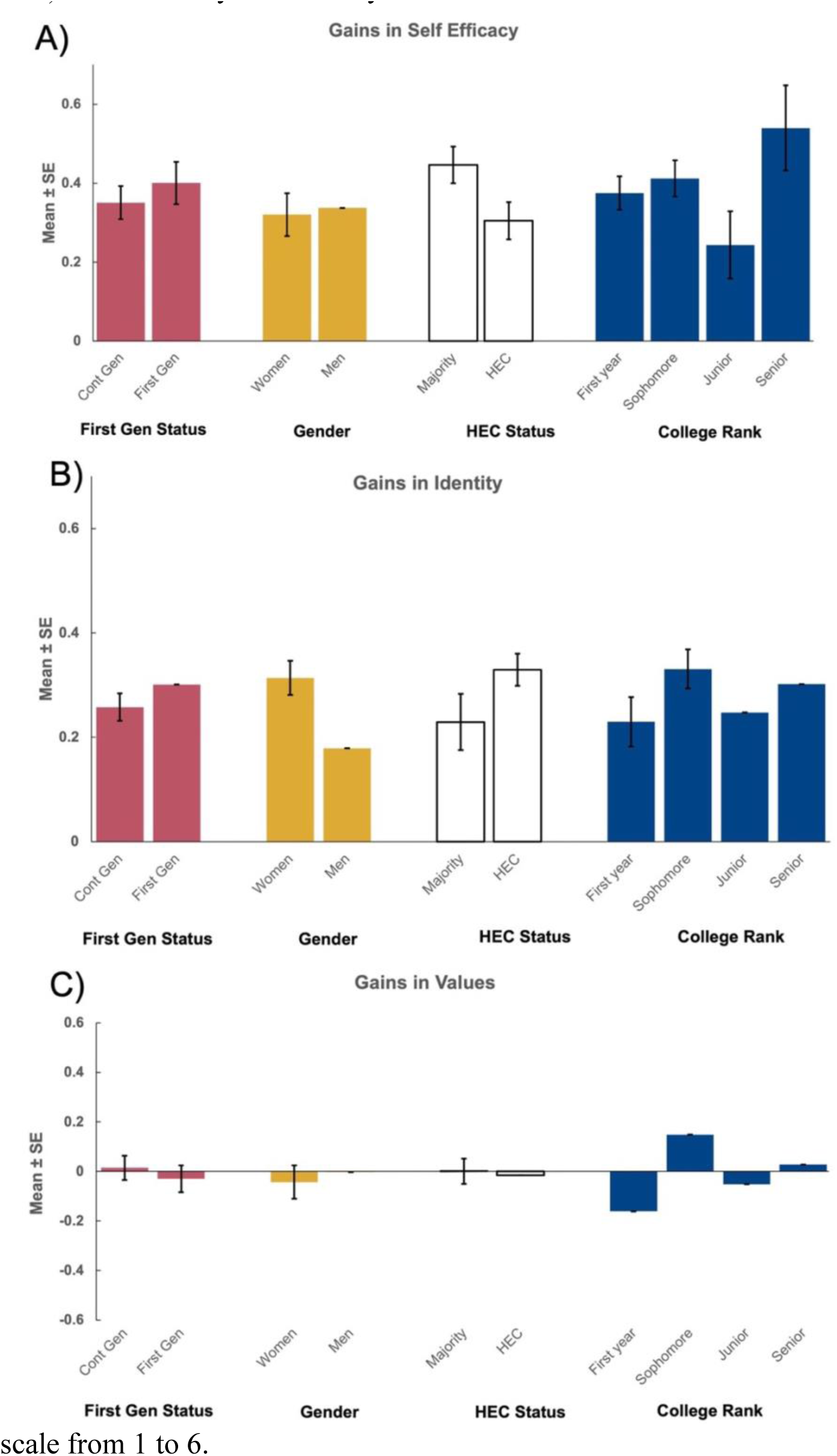
Mean student gains in (A) scientific self-efficacy, (B) scientific identity, and (C) orientation toward scientific values, differentiated by student demographics. (*N = 698; p < 0.05)*. Self-efficacy and identity were measured on scales from 1 to 5. Values were measured on a scale from 1 to 6.

Similarly, the analysis revealed that all demographic groups showed significant gains in *scientific identity,* and, partially consistent with our hypothesis, the analysis found no meaningful group-mean differences based on first-generation status or HEC status within classes. Inconsistent with our predictions, we found that women exhibited larger gains in their scientific identity compared to their male classmates (**Fig. 1B & Supplemental Tables)**.

Finally, the analysis revealed no meaningful group-mean differences in *scientific values* based on first-generation status, gender identity, or HEC status within-classes. However, the control variable of year in school revealed that sophomores had higher gains that first-year students (**Fig. 1C & Supplemental Tables)**.

#### Hypothesis 2b

*Tiny Earth positively affects TIMSI mechanisms across all course characteristics*. The analysis revealed that students in courses of all types exhibited significant gains in *scientific self-efficacy* and, partially consistent with our hypothesis, the analysis found no meaningful between-course differences based on course type, division, format, modality, the proportion of first-generation students in the class, or proportion of women in the class. Contrary to our hypothesis, high-enrollment courses exhibited larger course-average gains in self-efficacy than did small-and medium-enrollment courses (**Fig. 2A & Supplemental Tables**). Further, courses with lower proportions of students from HEC groups exhibited larger course-average gains in self-efficacy than did courses with higher proportions of students from HEC groups (**Fig. 2D & Supplemental Tables**).

**Fig. 2.**
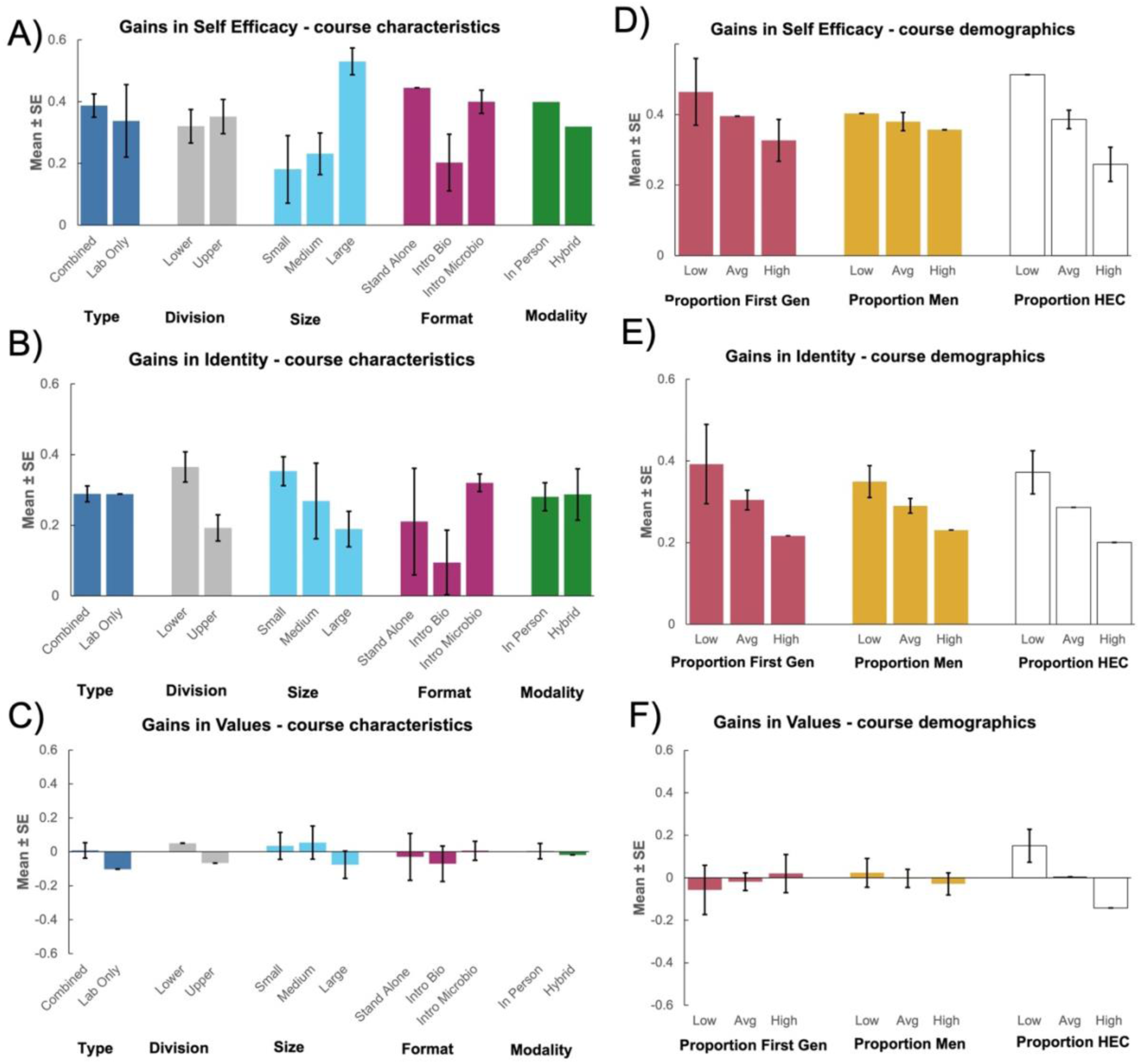
Mean student gains in (A) scientific self-efficacy, (B) scientific identity, and (C) orientation toward scientific values, differentiated by course characteristics and proportion of (D) first-generation students, (E) men, and (F) students from historically excluded communities (HEC) (*N = 698; p < 0.05*). Self-efficacy and identity were measured on scales from 1 to 5. Values were measured on a scale from 1 to 6.

Similarly, students in courses of all types exhibited significant positive gains in *scientific identity*, and partially consistent with our hypothesis, we found no meaningful between-course differences based on the type, size, modality, or the proportion of first-generation, women, or HEC students in the course. Inconsistent with our hypothesis, the analysis revealed that lower-division courses exhibited larger gains course-average *scientific identity* than did upper-division courses, and students enrolled in Tiny Earth curriculum that was integrated into microbiology courses exhibited larger gains than students in those integrated into biology courses (**Fig. 2B**, **Fig. 2E, & Supplemental Tables**).

Finally, the analysis indicated that there were no meaningful between-course differences in *scientific values* based on course type, division, size, format, modality, or the proportion of first-generation or women students in the course (**Figs. 2C & Supplemental Tables**). Inconsistent with our hypothesis, courses with lower proportions of students from HEC groups exhibited larger course-average gains in values than did courses with higher proportions of students from HEC groups (**Fig. 2F & Supplemental Tables**).

#### Hypothesis 3

*Tiny Earth positively affects persistence intentions.* A multilevel regression model indicated that the starting course-average level of scientific persistence intentions was moderately high (i.e., course average 62% of the maximum possible score). Inconsistent with our predictions, the analysis revealed course average gain in *persistence intentions* across classes was not significantly different from zero (**Table 3**).

#### Hypothesis 4a

*Tiny Earth positively affects persistence intentions for all student demographics*. Since a multilevel model revealed no between-course variability in persistence intention gain scores, multiple regression was used to address hypothesis 4a. Partially consistent with our predictions, the analysis revealed no meaningful group-mean differences in *persistence intentions* gain scores between women and men, between students from majority or HEC groups, or based on year in school. However, inconsistent with our predictions we found that first-generation students exhibited larger gains in persistence intentions than their continuing-generation peers (**Fig. 3 & Supplemental Tables**).

**Fig. 3.**
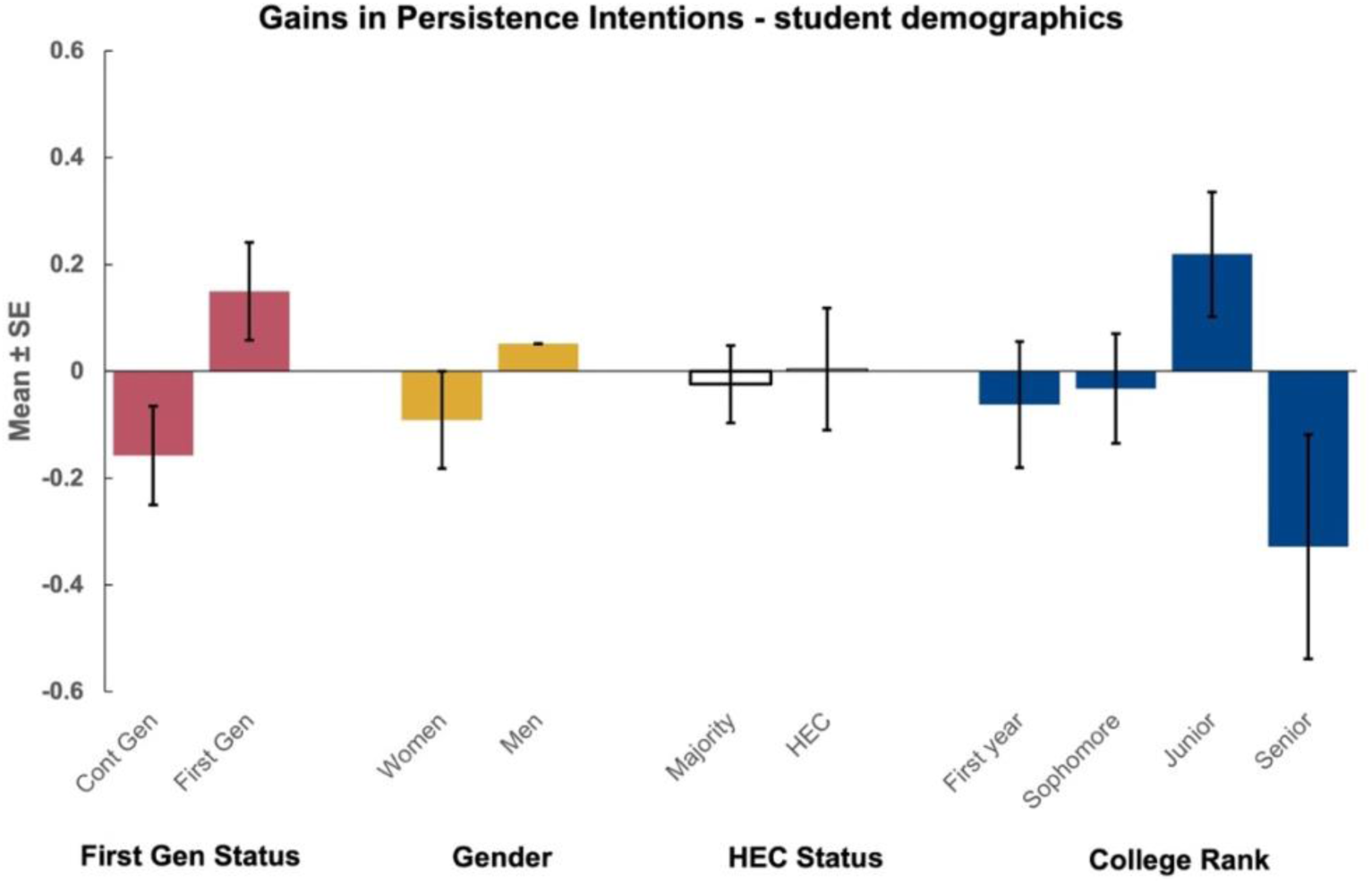
Mean student gains in intentions to persist in STEM, differentiated by student demographics. (*N = 698; p < 0.05*). Persistence was measured on a scale from 0 to 10.

#### Hypothesis 4b

*Tiny Earth positively affects persistence intentions for all course characteristics.* Consistent with our predictions, the analysis revealed no meaningful between-course differences based on course type, division, enrollment size, format, or modality **(Fig. 4 & Supplemental Tables)**.

**Fig. 4.**
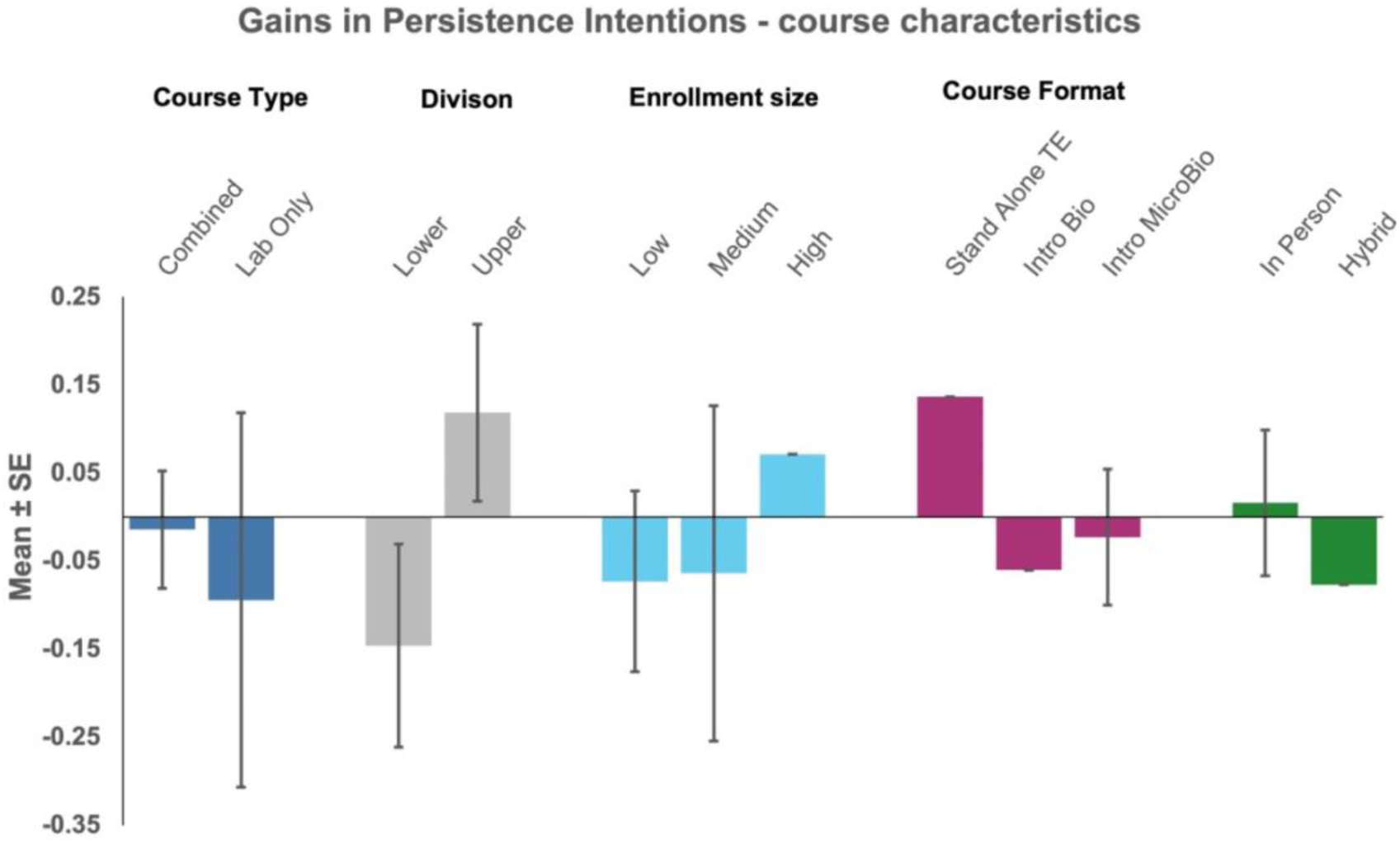
Mean student gains in intentions to persist in STEM, differentiated by course characteristics. (*N = 698; p < 0.05*). Persistence was measured on a scale from 0 to 10.

#### Hypothesis 4c

*Student gains in the three TIMSI mechanisms predict persistence intentions*. Consistent with the TIMSI theory, students’ gains in each of the TIMSI mechanisms (*scientific self-efficacy*, *identity*, and *values*) were positively associated with gains in *persistence intentions*. Students who reported the highest gains in scientific self-efficacy, identity, and values showed the highest gains in persistence intentions. Gains in scientific identity exhibited the largest influence, followed by gains in scientific values and self-efficacy (**Fig. 5**, **Table 4, & Supplemental Tables**).

**Fig 5.**
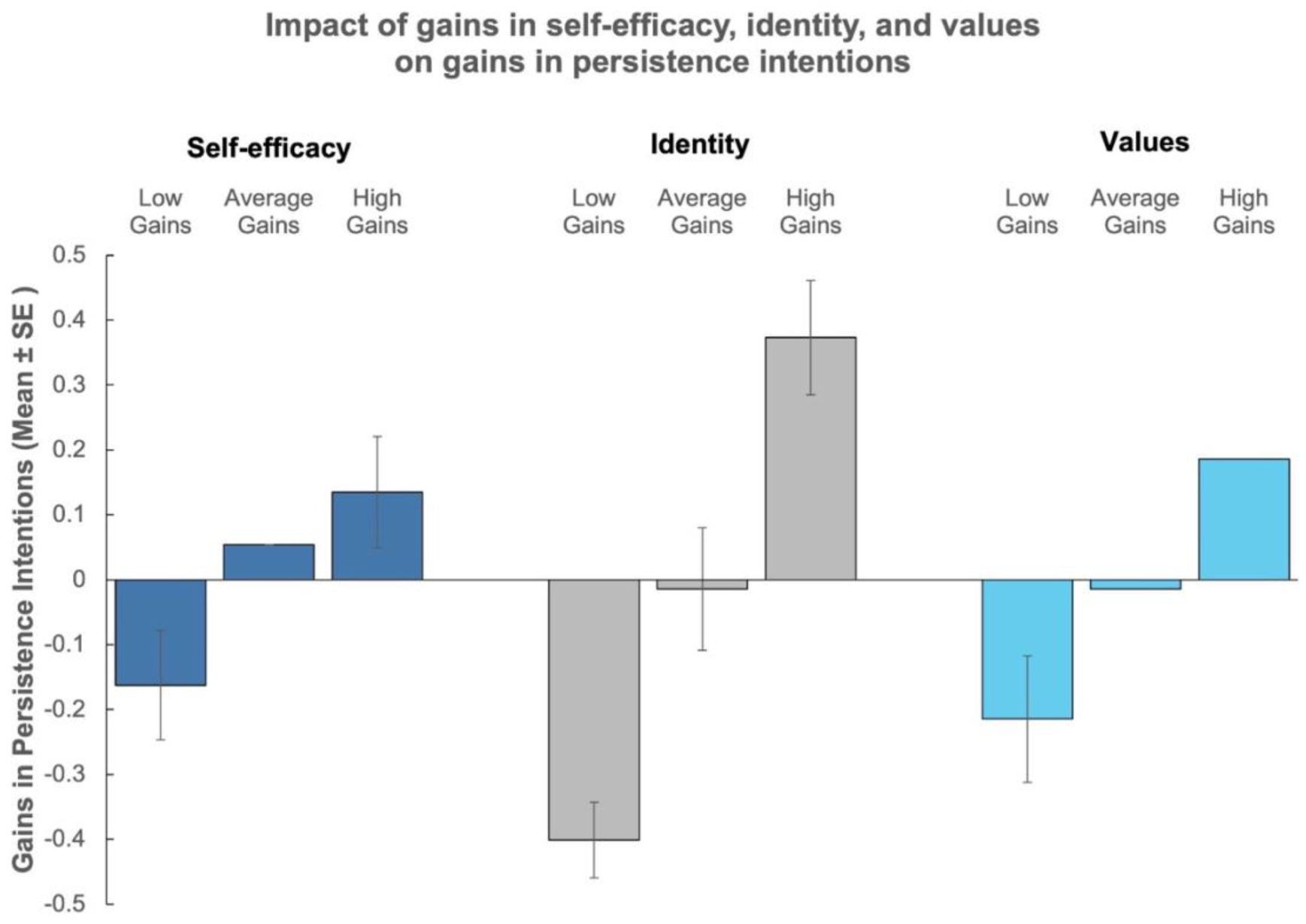
Relationship between gains in students’ scientific self-efficacy, identity, and values with gains in intent to persist in STEM. (*N = 698; p < 0.05*).

**Table 4.** Correlations between students’ scientific self-efficacy (E), identity (I), values (V), and persistence intentions (P); student demographics; and course characteristics (*N* = 698).

| Variable | 1 | 2 | 3 | 4 | 5 | 6 | 7 | 8 | 9 |
| --- | --- | --- | --- | --- | --- | --- | --- | --- | --- |
| **1. $\Delta$ Persistence Intentions** | 1 |  |  |  |  |  |  |  |  |
| **2. $\Delta$ Efficacy** | .19* | 1 |  |  |  |  |  |  |  |
| **3. $\Delta$ Identity** | .30* | .29* | 1 |  |  |  |  |  |  |
| **4. $\Delta$ Values** | .22* | .28* | .33* | 1 |  |  |  |  |  |
| **5. HEC Status<sup>a</sup>** | .00 | -.07 | .04 | -.03 | 1 |  |  |  |  |
| **6. Male Status<sup>b</sup>** | .00 | -.02 | -.11* | .00 | -.02 | 1 |  |  |  |
| **7. Trans/NB/GF Status<sup>b</sup>** | -.07 | .00 | -.01 | .02 | .04 | -.06 | 1 |  |  |
| **8. First-Gen Status<sup>c</sup>** | .04 | -.01 | .01 | -.04 | .36* | -.10* | .01 | 1 |  |
| **9. Sophomore<sup>d</sup>** | .00 | .01 | .04 | .11* | -.03 | -.04 | .01 | -.08* | 1 |
| **10. Junior<sup>d</sup>** | .08* | -.08* | -.03 | -.02 | -.03 | .09* | .00 | -.05 | -.41* |
| **11. Senior<sup>d</sup>** | -.06 | .04 | -.01 | .00 | .17* | .03 | .05 | .14* | -.25* |
| **12. Lab only<sup>a</sup>** | .00 | .01 | .03 | -.01 | -.17* | -.02 | -.04 | -.16* | .18* |
| **13. Upper Division<sup>b</sup>** | .06 | .02 | -.06 | -.04 | -.11* | .10* | -.03 | -.15* | -.08* |
| **14. Low Enrollment<sup>c</sup>** | -.01 | -.04 | .01 | -.01 | -.04 | -.05 | .05 | -.06 | -.15* |
| **15. Medium Enrollment<sup>c</sup>** | .04 | .01 | .03 | .03 | -.28* | .02 | -.01 | -.35* | .22* |
| **16. Hybrid<sup>d</sup>** | -.06 | -.07 | .02 | .01 | .17* | -.06 | -.07 | .25* | .03 |
| **17. Standalone Tiny Earth<sup>e</sup>** | .02 | .03 | .06 | .02 | -.13* | -.03 | -.03 | -.18* | .26* |
| **18. Integrated into Biology<sup>e</sup>** | -.03 | .03 | .02 | .02 | -.03 | -.09* | .03 | .00 | -.01 |

## DISCUSSION

This study aimed to better understand how an undergraduate science CURE with widespread implementation influences the processes leading to social integration among diverse students and classrooms. The study applied the Tripartite Integration Model of Social Influence (TIMSI) to understand how enrollment in the Tiny Earth CURE affects scientific self-efficacy, scientific identity, and orientation toward scientific values, and how these three attributes relate to intentions to persist in STEM. By examining individual student differences and determining how the class context impacts outcomes, **we established that the Tiny Earth CURE significantly increased student integration into the scientific community.** Students from both historically underrepresented and majority groups gained substantially in scientific self-efficacy and identity, which, along with values orientation, were linked to gains in intentions to persist in STEM. The findings demonstrate a robust effect of the Tiny Earth CURE across classroom contexts and student demographics.

As in previous studies of CUREs, we found that scientific self-efficacy increased across all student groups in the Tiny Earth CURE. Our study extends previous findings by examining how course characteristics impact outcomes and demonstrating that gains were slightly smaller for students in classes that had a higher proportion of HEC students and, for those in small-and medium-sized classes, and juniors. Course-level outcomes can be interpreted in several ways. While it may indicate that the Tiny Earth curriculum has differential outcomes because of the proportion of HEC students or the size of the classroom, it is equally possible that universities with higher proportions of HEC students are likely to be lower resourced and considered less competitive for entry. Data indicate that minority-serving institutions (MSIs) face significant underfunding while playing a significant role in bolstering the STEM workforce (National Academies of Sciences, Engineering, and Medicine, 2019).

We found that scientific identity increased across all groups of students and that the gains were greater among women, as others have also shown (Lorenzo et al., 2006). Scientific identity predicts persistence in STEM fields four years after graduation (Estrada, Hernandez, et al., 2018), indicating the importance of student maintaining their identification with science. Importantly, the Tiny Earth CURE appears to be interrupting the disidentification process that students, particularly HEC and first-generation students, experience in typical core science courses. Further research is needed to better understand how the antiracist, just, equitable, diverse, and inclusive (AJEDI) Tiny Earth content contributes to the increase in scientific identity among populations that have been historically excluded and marginalized in the scientific community.

An unanticipated outcome of this study was the high number of students who declined to identify their gender. Twenty-five percent of students selected ‘prefer not to say’ (PNTS) for the gender identity item, in contrast to <1% for race/ethnicity and academic rank. The PNTS item is different than the ‘other’ item on the instrument, from which we were able to identify a subset of students who specified their identity as transgender, non-binary, or gender-fluid. Although the number of PNTS is substantial, we cannot interpret what PNTS means in this context and therefore cannot aggregate them into a group for meaningful comparison. As a research community, we need to explore better ways for respondents to claim their identity on their own terms and ensure their contributions are not omitted from research.

Students are oriented toward scientific values when they internalize and value scientific experiences, such as building new knowledge to solve global challenges, the thrill of discovery, and the importance of discourse. In this study, orientation toward scientific values did not increase across all groups, but some groups showed significant increases. For example, values growth was greater for students in smaller classes than large ones, and sophomores grew more than their counterparts.

Perhaps the most dramatic finding in this study was the discovery that, consistent with the TIMSI model, changes in intention to persist in STEM were primarily explained by gains in scientific self-efficacy, identity, and values (Figure 5). The striking outcome is that those who experienced gains in any of the three elements were far more likely to experience gains in the intent to persist in STEM. A second important finding was that first-generation students experience larger gains in their persistence intentions than their continuing-generation peers. This study therefore provides further quantitative support for the powerful TIMSI model as a predicter of persistence in STEM.

The findings reported here provide support for the recommendations of Handelsman et. al. (2022) that achieving STEM diversity requires reforming teaching practices, creating welcoming classrooms and expanding relevance to diverse groups, which the Tiny Earth CURE explicitly aims to achieve. Although many STEM educators hope that their courses will inspire students to remain in STEM, in reality, the core science courses commonly discourage college students from continuing in science. Around 60% of those students intending to major in science at the start of college seek non-STEM majors after taking the introductory curriculum. Data show that the exodus from science is particularly pronounced for students who are first generation in college or from marginalized communities, making the current system discriminatory against certain groups. Some college-level STEM educators would rejoice at these statistics, assuming that they provide evidence that their courses weed out students not fit for the rigors of science. But in fact, the learning environment in many introductory courses repels numerous students who receive high grades, indicating that the introductory curriculum discourages students who could be significant contributors to science. Therefore, programs that increase, or simply prevent a decline in, student intentions to persist are essential to enlarging the cohort of STEM college graduates (Schultz et al., 2011).

Rather than *hoping* to inspire students, colleges and universities need to confront the data. Higher education should be employing interventions, such as CUREs, that reliably counter the steady loss of good students from STEM. One research group estimated that if every undergraduate in the United States could take a CURE, an estimated 200,000 more students would graduate college in STEM fields each year (Dolan & Weaver, 2021). Considering that several CUREs have been rigorously studied, optimized, and disseminated, there are diverse, validated choices. National policy should provide resources for training to teach a CURE for one instructor from each of the 4,000 undergraduate institutions in the United States. We estimate that such training could be accomplished for $25 million and eventually could result in every undergraduate having the option to take a research course. This seems like a modest investment to exercise a powerful lever that could create a sufficient, diverse, and robust workforce for science and technology fields.

## ACCESSING MATERIALS

The Tiny Earth student research guide is available for purchase online at XanEdu Publishing, which includes a print book plus FlexEd digital course access at https://www.xanedu.com/catalog-product-details/tiny-earth. Instructors may request a free desk copy directly from the publisher. Some materials are available in previous publications (e.g., Hernandez et al., 2018, González-Orta et al., 2022, Miller et al., 2022). To access the full set of instructor resources, apply to attend a TEPI Training at https://tinyearth.wisc.edu/tiny-earth-partner-instructors/. Trainings are currently free for US college instructors, except for travel to the training site.

## ACKNOWLEDGMENTS

Work reported in this publication was supported by the National Institutes of Health Common Fund and Office of Scientific Workforce Diversity under award U54 GM119023 (NRMN), administered by the National Institute of General Medical Sciences. UCSF IRB #19-28867.

We thank the many Tiny Earth instructors, students, and staff who made this study possible. The Tiny Earth Partner Instructor (TEPI) Training, mentor pairing, and pivot to online was made possible by Dr. Debra Davis; Trang Tran; Martel DenHartog; staff and students at Johns Hopkins University, University of Connecticut, and University of Wisconsin-Madison; and many TEPI facilitators. The online components of the Tiny Earth curriculum were made possible by Dr. Enid González-Orta, Dr. Aarti Raja, and many other TEPIs.

We dedicate this paper to all the stalwart TEPIs and Tiny Earthlings (the students) who persevered in advancing scientific research and education during the COVID-19 pandemic.

### Box. 1. Student Learning Objectives for the Tiny Earth Course.

**Module 1: Isolating Bacteria from Soil**

➢ Apply aseptic and serial dilution plating techniques to obtain isolated bacterial colonies from soil.
➢ Develop hypotheses about how bacteria from various soil environments vary in growth, diversity, and abundance on various laboratory media.
➢ Design experiments, gather and analyze data, and summarize findings.
➢ Estimate the number of microorganisms in a sample.

**Module 2: Screening Bacterial Isolates for Antibiotic Production**

➢ Screen bacterial isolates for antimicrobial activity against a panel of safe relatives of known pathogens or other environmental microbes.
➢ Develop hypotheses about bacterial interactions and antibiotic production using various media.
➢ Design experiments, gather and analyze data, and summarize findings.
➢ Explain how bacteria harness nutrients from their environment to grow.
➢ Explain the importance of studying clinically relevant microbes, distinguishing between pathogenic, attenuated, and non-pathogenic microbes.
➢ Propose narrow-spectrum antibiotics by leveraging the differences between Gram-positive and Gram-negative bacteria.

**Module 3: Characterizing the Antibiotic-producing Bacterial Isolates**

➢ Integrate molecular, morphological, and biochemical characterization methods to establish a putative taxonomic identity for one or more environmental bacterial isolates.
➢ Analyze and interpret biochemical test data to infer physiological, physical, and/or pathogenic properties of isolates.
➢ Justify why the 16S rRNA gene is a suitable molecular target to group and identify bacteria.
➢ Explain how taxonomic identity can be useful for prioritizing antibiotic-producing bacterial isolates for future study.

**Module 4: Advancing the Antibiotic Discovery Pipeline**

➢ Extract metabolic compounds from bacterial isolates and design experiments to test the compounds for antimicrobial activity.
➢ Gather, visualize, and interpret data; communicate findings.
➢ Explain the processes of antibiotic discovery and resistance development in pathogen populations.
➢ Propose a candidate antibiotic that would be effective against bacteria but safe for humans; explain how the antibiotic would target bacterial functions not found in humans.

### Box. 2. Sample Tiny Earth Partner Instructor (TEPI) Training Schedule.

➢ **Pre-Training:** Interactive Welcome Tutorial
➢ **Monday:** Welcome, Overview of Tiny Earth, and Lab 1 (Isolating Bacteria from Soil)
➢ **Tuesday:** Scientific Teaching and Lab 2 (Screening Bacterial Isolates for Antibiotic Production)
➢ **Wednesday:** Inclusive Teaching and Lab 3 (Characterizing the Antibiotic-producing Bacterial Isolates)
➢ **Thursday:** Online Adaptations, Tiny Earth Database, and Lab 4 (Advancing the Antibiotic Discovery Pipeline)
➢ **Friday:** Group Presentations and Lessons from Veteran TEPIs
➢ **Post-Training:** Mentor Pairings, Network Activities, and Teaching Tiny Earth

