## Supplemental Materials and Tables for "Tiny Earth CURE improves student persistence in science"

This document contains information about preliminary analyses to establish evidence of measurement validity across time and between groups, specifically:

- the longitudinal measurement invariance of the TIMSI processes and outcome used at pre- and post-test in this study (Table S1); and
- the cross-group (gender and race/ethnicity) measurement invariance of the TIMSI processes and outcome used at pre- and post-test in this study (Table S2).

### Analytical Approach and Preliminary Analyses

The preliminary statistical analyses were conducted in three stages.

**Stage 1** included preliminary analysis of statistical assumption checks and missing data analysis to understand whether our statistical models would produce accurate and unbiased statistical results. Missing data that are not consistent with missing completely at random can introduce bias into statistical model results (R. J. A. Little, 1988). Therefore, we first tested the assumption that the pattern of missing student-level data outcomes were consistent with missing completely at random (MCAR) via Little's MCAR test (R. J. A. Little, 1988) in the overall data set. The results indicated missing data in the outcomes were consistent with the MCAR assumption ($\chi^2$ [$df = 6$] = 5.87, $p = 0.47$). Similarly, outliers in data can bias the results of statistical analyses. Therefore, we then conducted outlier analysis using leverage values (i.e., high leverage values indicate cases that have extreme scores on a set of predictors), studentized deleted residuals (SDRs; i.e., large SDRs indicate cases that have extreme outcome scores, given their pattern on the predictors) and Cook's distance (i.e., large Cook's distance values indicate cases that have both extreme values on the predictors and outcome) measures and found no evidence of extreme outliers (Judd et al., 2017). Next, we tested distributional assumptions of the outcomes and found that the data were not normally distributed, but the variances on the outcomes were consistent across groups (i.e., homoscedastic), and had linear relationships with the predictors (Judd et al., 2017). Taken together, these findings indicate that the data meet the assumptions required to produce accurate and unbiased results in the statistical analyses.

**Stage 2** included testing the longitudinal measurement properties of outcomes to determine whether the outcomes exhibited consistent measurement properties over time, which is an essential characteristic for assessing unbiased change in outcomes over time. We assessed the measurement properties of each of the outcomes based on comparison of pre- and post-course measures using a series of confirmatory factor analysis (CFA) models (Kline, 2016), which tests the longitudinal measurement invariance of each outcome over time (T. D. Little, 2013). These tests determine whether the measurement properties of the outcomes are consistent over time. For each longitudinal measurement invariance test, we compared three invariance models: configural (constant factor structure across time), metric (constant factor loadings across time), and scalar (constant indicator intercepts across time). Together, these models test for consistency in how the latent outcome(s) related to student responses to the questions over-time and consistency in the average observed response to each question over-time.

In addition to reporting the global chi-square test, we evaluated global data-model fit with the following fit indices and cut-off values (Kline, 2016): the comparative fit index (CFI; values $\geq$ 0.95 indicating acceptable fit), root mean square error of approximation (RMSEA; values $\leq$ 0.08 or 90% CI that included 0.05 but not 0.10 indicating acceptable fit), and the standardized root mean square residual (SRMR; values $\leq$ .08 indicating acceptable fit). When comparing CFA models for invariance tests, we used the following indices to indicate poor model fit: changes in CFI > 0.010 and changes in RMSEA > 0.015 (Chen, 2007; Cheung & Rensvold, 2002).

The results of the longitudinal measurement invariance tests revealed that scientific self-efficacy, identity, values, and career persistence intentions fit the data well from pre- and post-course surveys, and that measurement properties were consistent over time (i.e., longitudinal scalar invariance holds (**Table S1**).

***Longitudinal measurement invariance tests.*** As noted above, in stage 2 of the preliminary analyses, we tested the longitudinal measurement invariance of the TIMSI processes (i.e., scientific self-efficacy, identity, and community values), as well as the outcome of science career persistence intentions. Longitudinal measurement invariance tests were conducted separately for each of the four constructs within the entire analytic sample ($N$ = 698). The tests involved comparing three nested models. First, as a configural invariance model that specified the same factor structure in which all indicators measured at Time-1 loaded only on the latent factor at Time-1 (e.g., Scientific Self-Efficacy T1) and similarly the indicators at Time-2 loaded on the latent factor onto the factor at Time-2, but the factor loadings and indicator intercepts were freely estimated. Further, as is customary for longitudinal models, the configural model allowed auto-correlated residuals (e.g., residual of indicator 1 at Time-1 allowed to correlate with the residual of indicator 1 at Time-2; Little, 2013). Second, the metric invariance involved constraining the factor loadings of the indicators to be the same at Time-1 and Time-2 (e.g., constraining the loading of indicator 1 at Time-1 to be the same as indicator 1 at Time-2). Third, the scalar invariance model involved constraining the indicator intercepts to be the same at Time-1 and Time-2 (e.g., constraining the intercept of indicator 1 at Time-1 to be the same as the intercept of indicator 1 at Time-2), while constraining the latent mean of the construct at Time-1 to zero and allowing the Time-2 latent mean of the construct to be freely estimated. This structure allows the latent mean at Time-2 to change (e.g., grow), while holding the measurement model constant.

After estimating each of the three models, global fit was assessed using the criteria outlined in the main narrative. If global fit indices indicated potential problems in the data-model fit, we examined local fit statistics (i.e., standardized residual means and covariances) to identify potential sources of mis-fit. Re-specification, such as adding additional residual covariances, involved carefully evaluating the wording of individual items along with the local fit indices.

As shown in Table S1, we fit longitudinal measurement invariance models to the Science Career Persistence Intentions scale at pre- and post-test. The configural model (M1.1) indicated relatively poor model fit. A close inspection of local fit indices and item wording revealed that word question formatting introduced correlations in responses to some questions over and above the relationship between the individual items and the latent construct. Specifically, indicators 1, 2, and 3 exhibited residual correlations at each timepoint (1. “To what degree do you plan to pursue a biomedical science-related research career?,” 2. “To what degree do you plan to obtain a biomedical science-related undergraduate degree (e.g., B.A. or B.S.)?," 3. "To what degree do you plan to obtain a biomedical science-related graduate degree (e.g., Masters or PhD)?"), as well as a residual correlation between the indicator 2 and 7 (7. "To what degree do you plan to pursue a career in which you will present scientific papers at conferences?"). Although added complexity of the residual matrix is not ideal, we modeled the additional residual covariances to fully capture the complexity, which substantially improved model fit and resulted in an acceptably fitting model (i.e., M1.2). Constraining the factor loadings to be constant over time (i.e., M1.3) and constraining indicator intercepts to be constant over time (i.e., M1.4) did not worsen model fit.

Concerning scientific self-efficacy, the longitudinal measurement invariance tests revealed that the configural invariance model provided acceptable fit for the data (M2.1). Further, constraining the factor loadings (M2.2) and the indicator intercepts (M2.3) did not worsen model fit.

Concerning scientific identity, the longitudinal measurement invariance tests revealed that the configural invariance model provided marginally acceptable fit to the data (M2.1). A close inspection of local fit indices and item wording revealed that word question formatting introduced correlations in responses to some questions. Specifically, indicators 1 and 3 exhibited residual correlations at each timepoint (1. "I have a strong sense of belonging to the community of scientists." and 3. "I feel like I belong in the field of science."). As above, we modeled the additional residual covariances to fully capture the complexity, which substantially improved model fit and resulted in an acceptably fitting model (M3.2). Constraining the factor loadings to be constant over time (i.e., M3.3) did not worsen model fit. Constraining the indicator intercepts resulted in poorer fit (M3.4); however, a close inspection of the local fit indices revealed that only indicator 2 exhibited non-constant intercepts over-time. Therefore, we relaxed the intercept constraint for indicator 2 and achieved partial scalar invariance, which did not worsen model fit (M3.5).

Concerning scientific values, the longitudinal measurement invariance tests revealed that the configural invariance model provided acceptable fit for the data (M4.1). Further, constraining the factor loadings (M4.2) and the indicator intercepts (M4.3) did not worsen model fit.
Stage 3 involved testing the cross-group measurement invariance of all outcomes at each time point as a function of gender (female, male, other genders/did not report) and race and ethnicity (HEC groups, racial and ethnic majority) to determine whether the outcomes exhibited consistent measurement properties across demographic groups. Cross-group measurement invariance is an essential characteristic for assessing unbiased differences across groups. As with longitudinal-measurement invariance, we compared the groups across three model tests (configural, metric, and scalar invariance) and used the same global-model fit statistics and criteria to determine acceptable fit. The cross-group measurement invariance tests showed that the outcomes fit the data equally well across demographic groups at pre- and post-course (**Table S2**).

Overall, we conclude that these tests show robust evidence of longitudinal measurement invariance for each construct over-time.

**Stage 3** involved testing the cross-group measurement invariance of all outcomes at each time point as a function of gender (female, male, other genders/did not report) and race and ethnicity (HEC groups, racial and ethnic majority) to determine whether the outcomes exhibited consistent measurement properties across demographic groups. Cross-group measurement invariance is an essential characteristic for assessing unbiased differences across groups. As with longitudinal-measurement invariance, we compared the groups across three model tests (configural, metric, and scalar invariance) and used the same global-model fit statistics and criteria to determine acceptable fit. The cross-group measurement invariance tests showed that the outcomes fit the data equally well across demographic groups at pre- and post-course (**Table S2**).

***Cross-group measurement invariance tests.*** As noted above, we tested the cross-group measurement invariance of the TIMSI processes and science career persistence intentions in stage 3 of the preliminary analyses. Cross-group measurement invariance tests were conducted across all latent constructs (e.g., scientific self-efficacy, identity, values, and persistence intentions) separately for Time-1 and Time-2. For gender, we compared the following three groups: females, males, and other genders. The other gender category included a small number of individuals that self-identified as transgender or non-binary, as well as those that preferred to not report their gender identity. For race/ethnicity, we compared students from historically excluded communities (HEC) and those from historically included communities (i.e., White and Asian). Sample sizes for this groups are provided in the main text Table 1. The cross-group measurement invariance tests involved comparing three nested models. First, as a configural invariance model that specified the same factor structure for all groups, but the factor loadings and indicator intercepts were freely estimated. In addition, complexities in the residual covariance matrix found for the longitudinal measurement models were replicated in the cross-group tests. Second, the metric invariance involved constraining the factor loadings of the indicators to be the same across all groups (e.g., constraining the loading of Scientific Self-Efficacy indicator 1 to be the same in the URM and Majority groups). Third, the scalar invariance model involved constraining the indicator intercepts to be the same across all groups (e.g., constraining the intercept of Scientific Self-Efficacy indicator 1 to be the same in the URM and Majority groups). After estimating each of the three models, global fit was assessed using the criteria outlined in the main narrative and potential problems with data-model fit, as well as the approach to re-specification were the same as described in the paragraphs above.

As shown in Table S2, when testing gender-based cross-group measurement invariance the configural model provided acceptable fit to the data at both Time-1 (M5.1) and Time-2 (M6.1). Further, constraining the factor loadings at each time-point (M5.2 & M6.2) and the indicator intercepts at each time-point (M5.3 & M6.3) did not worsen model fit. We conclude that these tests show robust evidence of gender-based cross-group measurement invariance for each construct at each time-point.

Similarly, when testing HEC-based cross-group measurement invariance the configural model provided acceptable fit to the data at both Time-1 (M7.1) and Time-2 (M8.1). Further, constraining the factor loadings at each time-point (M7.2 & M8.2) and the indicator intercepts at each time-point (M7.3 & M8.3) did not worsen model fit. We conclude that these tests show robust evidence of race-based cross-group measurement invariance for each construct at each time-point.

**SUPPLEMENTAL TABLES**
**Table S1. Correlations between course characteristics (Level-2; J = 31).**

| Variable | **1** | **2** | **3** | **4** | **5** | **6** | **7** | **8** | **9** | **10** |
|---|---|---|---|---|---|---|---|---|---|---|
| 1. Lab Only[a] | 1 | | | | | | | | | |
| 2. Upper Division[b] | -.12 | 1 | | | | | | | | |
| 3. Low Enrollment[c] | .05 | -.19 | 1 | | | | | | | |
| 4. Medium Enrollment[c] | .28 | -.17 | .23 | 1 | | | | | | |
| 5. Hybrid[d] | -.11 | .21 | -.09 | -.77* | 1 | | | | | |
| 6. Standalone Tiny Earth[e] | .01 | -.13 | -.02 | .07 | -.17 | 1 | | | | |
| 7. Integrated into Biology[e] | .49* | -.08 | .07 | .13 | -.02 | .07 | 1 | | | |
| 8. Avg. First-gen Status | .05 | -.09 | -.34 | .05 | -.09 | .01 | -.21 | 1 | | |
| 9. Avg. HEC Status | .10 | -.22 | -.34 | -.01 | -.30 | .16 | -.18 | -.05 | 1 | |
| 10. Avg. Male Status | .00 | -.03 | -.13 | -.07 | .07 | -.19 | -.15 | -.19 | .15 | 1 |

Notes: Combined Course: Lab-plus-lecture format. Low-enrollment class: 0-19 students; medium-enrollment class: 20-100; high-enrollment class: more than 100. [a]Reference group = combined mode (lab and lecture). [b]Reference group = lower-division course. [c]Reference group = high-enrollment class. [d]Reference group = in-person modality. [e]Reference group = integrated into microbiology course. *$p < 0.05$.

**Table S2. Student Level Descriptive Statistics (N=698)**

| Variable | M | SD | Minimum Value | Maximum Value | Skew | Kurtosis |
|---|---|---|---|---|---|---|
| Persistence Intentions T1 | 6.11 | 2.13 | 0.00 | 10.00 | -0.29 | -0.51 |
| Scientific Self-Efficacy T1 | 3.37 | 0.84 | 1.00 | 5.00 | -0.47 | 0.41 |
| Scientific Identity T1 | 3.26 | 0.97 | 1.00 | 5.00 | -0.21 | -0.48 |
| Scientific Values T1 | 4.97 | 0.96 | 1.00 | 6.00 | -1.09 | 1.11 |
| Persistence Intentions T2 | 6.10 | 2.21 | 0.00 | 10.00 | -0.38 | 2.64 |
| Scientific Self-Efficacy T2 | 3.74 | 0.85 | 1.00 | 5.00 | -0.66 | 3.64 |
| Scientific Identity T2 | 3.54 | 0.95 | 1.00 | 5.00 | -0.33 | 2.63 |
| Scientific Values T2 | 4.96 | 0.97 | 1.00 | 6.00 | -1.07 | 3.96 |
| ∆ Persistence Intentions | -0.01 | 1.63 | -8.71 | 7.00 | -0.40 | 2.16 |
| ∆ Efficacy | 0.37 | 0.87 | -4.00 | 4.00 | -0.19 | 1.79 |
| ∆ Identity | 0.27 | 0.78 | -4.00 | 3.00 | -0.32 | 2.26 |
| ∆ Values | -0.01 | 0.87 | -4.00 | 5.00 | -0.20 | 4.31 |
| HEC Status | 0.33 | 0.47 | 0.00 | 1.00 | 0.72 | -1.49 |
| Male Status | 0.22 | 0.41 | 0.00 | 1.00 | 1.34 | -0.20 |
| First-gen Status | 0.47 | 0.49 | 0.00 | 1.00 | 0.14 | -1.99 |
| Sophomore | 0.35 | 0.47 | 0.00 | 1.00 | 0.64 | -1.60 |

| | | | | | | |
|---|---|---|---|---|---|---|
| Junior | 0.24 | 0.43 | 0.00 | 1.00 | 1.20 | -0.57 |
| Senior | 0.10 | 0.30 | 0.00 | 1.00 | 2.62 | 4.85 |

Notes: N = sample size; M = mean; SD = standard deviation; Δ = gain scores between time 1 (T1) and time 2 (T2).

**Table S3. Summary of parameter estimates of multilevel models predicting TIMSI indicators of social integration**

| | Δ Scientific Self-efficacy | | | Δ Scientific Identity | | |
|---|---|---|---|---|---|---|
| Source | b | SE | 95% CI | b | SE | 95% CI |
| 1. Intercepts | 0.73 | 0.19 | [0.35, 1.11] | 0.23 | 0.19 | [-0.15, 0.60] |
| 2. First-gen Status[abd] | 0.05 | 0.08 | [-0.11, 0.21] | 0.04 | 0.05 | [-0.06, 0.15] |
| 3. HEC Status[abe] | -0.14 | 0.08 | [-0.30, 0.02] | 0.10 | 0.08 | [-0.06, 0.26] |
| 4. Male Status [abf] | 0.02 | 0.09 | [-0.16, 0.19] | -0.13 | 0.04 | [-0.22, -0.05] |
| 5. Trans/NB/GF[abf] | 0.08 | 0.18 | [-0.27, 0.44] | -0.11 | 0.15 | [-0.39, 0.18] |
| 6. Gender DNR[abf] | 0.18 | 0.11 | [-0.04, 0.40] | 0.14 | 0.04 | [0.07, 0.21] |
| 7. Sophomore[abg] | 0.04 | 0.06 | [-0.08, 0.16] | 0.10 | 0.07 | [-0.03, 0.24] |
| 8. Junior[abg] | -0.13 | 0.10 | [-0.33, 0.07] | 0.02 | 0.05 | [-0.08, 0.12] |
| 9. Senior[abg] | 0.16 | 0.10 | [-0.04, 0.37] | 0.07 | 0.18 | [-0.28, 0.43] |
| 10. Year DNR[abg] | 0.98 | 0.06 | [0.86, 1.10] | 0.96 | 0.06 | [0.83, 1.08] |
| 11. Lab Only[ah] | -0.05 | 0.14 | [-0.33, 0.23] | 0.00 | 0.09 | [-0.18, 0.18] |
| 12. Type DNR[ah] | -0.17 | 0.16 | [-0.48, 0.15] | -0.29 | 0.09 | [-0.47, -0.11] |
| 13. Upper Division[ai] | -0.05 | 0.10 | [-0.24, 0.14] | -0.17 | 0.07 | [-0.31, -0.03] |
| 14. Low-enrollment[aj] | -0.35 | 0.13 | [-0.61, -0.09] | -0.08 | 0.13 | [-0.34, 0.17] |
| 15. Medium-enrollment[aj] | -0.30 | 0.10 | [-0.49, -0.11] | -0.16 | 0.08 | [-0.336, 0.002] |
| 16. Stand Alone | 0.05 | 0.18 | [-0.31, 0.40] | -0.11 | 0.16 | [-0.42, 0.2] |
| 17. Biology | -0.20 | 0.11 | [-0.42, 0.02] | -0.23 | 0.11 | [-0.43, -0.02] |
| 18. Hybrid[ak] | -0.08 | 0.12 | [-0.32, 0.16] | 0.01 | 0.11 | [-0.2, 0.22] |
| 19. Format DNR[ak] | 0.10 | 0.23 | [-0.35, 0.54] | -0.58 | 0.14 | [-0.86, -0.3] |
| 20. Avg. First-gen Status[c] | -0.28 | 0.29 | [-0.85, 0.30] | -0.35 | 0.31 | [-0.96, 0.26] |
| 21. Avg. Male Status[c] | -0.26 | 0.33 | [-0.90, 0.38] | -0.66 | 0.34 | [-1.33, 0.01] |
| 22. Avg. HEC Status[c] | -0.67 | 0.29 | [-1.24, -0.09] | -0.45 | 0.23 | [-0.902, 0.003] |
| Level 1 $R^2$ | 0.017 | | | 0.022 | | |
| Level 2 $R^2$ | 1.000 | | | 1.000 | | |
| Total $R^2$ | 0.039 | | | 0.045 | | |

Table Continues…

Table Continued…

| | Δ Scientific Values | | |
|---|---|---|---|
| Source | b | SE | 95% CI |
| 1. Intercepts | -0.29 | 0.21 | [-0.71, 0.12] |
| 2. First-gen Status[abd] | -0.04 | 0.06 | [-0.17, 0.08] |
| 3. HEC Status[abe] | -0.02 | 0.06 | [-0.14, 0.11] |
| 4. Male Status [abf] | 0.04 | 0.10 | [-0.16, 0.25] |
| 5. Trans/NB/GF[abf] | 0.21 | 0.31 | [-0.39, 0.81] |
| 6. Gender DNR[abf] | 0.05 | 0.14 | [-0.21, 0.32] |
| 7. Sophomore[abg] | 0.31 | 0.09 | [0.14, 0.48] |
| 8. Junior[abg] | 0.11 | 0.10 | [-0.08, 0.30] |
| 9. Senior[abg] | 0.19 | 0.14 | [-0.09, 0.47] |
| 10. Year DNR[abg] | -0.03 | 0.10 | [-0.22, 0.16] |
| 11. Lab Only[ah] | -0.11 | 0.10 | [-0.31, 0.09] |
| 12. Type DNR[ah] | -0.05 | 0.14 | [-0.33, 0.23] |
| 13. Upper Division[ai] | -0.12 | 0.06 | [-0.24, 0.00] |
| 14. Low-enrollment[aj] | 0.02 | 0.14 | [-0.26, 0.30] |
| 15. Medium-enrollment[aj] | -0.11 | 0.13 | [-0.37, 0.15] |
| 16. Stand Alone[ak] | -0.04 | 0.15 | [-0.33, 0.26] |
| 17. Biology[ak] | -0.08 | 0.14 | [-0.35, 0.20] |
| 18. Hybrid[al] | -0.02 | 0.12 | [-0.26, 0.21] |
| 19. Modality DNR[al] | -0.87 | 0.20 | [-1.27, -0.47] |
| 20. Avg. First-gen Status[c] | 0.15 | 0.38 | [-0.59, 0.90] |
| 21. Avg. Male Status[c] | -0.29 | 0.47 | [-1.22, 0.64] |
| 22. Avg. HEC Status[c] | -0.77 | 0.29 | [-1.35, -0.19] |
| Level 1 $R^2$ | 0.017 | | |
| Level 2 $R^2$ | 0.719 | | |
| Total $R^2$ | 0.039 | | |

Notes: Only fixed effects of predictors were tested in this study. Statistical estimates indicate the effects of predictors on three outcomes for the focal group when compared to the reference group. First-gen = first-generation college student status (-.5 = no, .5 = yes). DNR = did not report / prefer not to say. [a]Effect coded such that the reference group = -0.5, and the focal group = 0.5. [b]Group-mean centered variable. [c]Grand-mean centered variable. [d]Reference group = continuing-generation college student. Reference group = racial majority student. [f]Reference group = female. [g]Reference group = first-year student. [h]Reference group = combined mode (lab and lecture). [i]Reference group = lower division. [j]Reference group = high-enrollment course. [k]Reference group = integrated into microbiology course. [l]Reference group = in-person.

**Table S4. Summary of Parameter Estimates of Regression Model Predicting Gains in Persistence Intentions ($F_{[21, 676]} = 4.87$, $p < .001$, $R^2 = .15$)**

| Source | b | SE | 95% CI |
|---|---|---|---|
| 1. Intercepts | -1.38 | 0.59 | [-2.53, -0.23] |
| 2. First-gen Status[abd] | 0.31 | 0.14 | [0.03, 0.59] |
| 3. HEC Status[abe] | 0.03 | 0.14 | [-0.25, 0.31] |
| 4. Male Status[abf] | 0.14 | 0.15 | [-0.15, 0.44] |
| 5. Trans/NB/GF[abf] | -1.03 | 0.48 | [-1.97, -0.09] |
| 6. Gender DNR[abf] | 0.23 | 0.17 | [-0.12, 0.57] |
| 7. Sophomore[abg] | 0.03 | 0.16 | [-0.28, 0.34] |
| 8. Junior[abg] | 0.28 | 0.17 | [-0.05, 0.61] |
| 9. Senior[abg] | -0.27 | 0.25 | [-0.75, 0.22] |
| 10. Year DNR [abg] | -1.30 | 0.61 | [-2.49, -0.11] |
| 11. Δ Scientific Self-efficacy | 0.17 | 0.07 | [0.03, 0.31] |
| 12. Δ Scientific Identity | 0.50 | 0.10 | [0.30, 0.69] |
| 13. Δ Scientific Values | 0.23 | 0.08 | [0.07, 0.39] |
| 14. Lab Only[ah] | -0.08 | 0.23 | [-0.54, 0.38] |
| 15. Type DNR [ah] | 0.29 | 0.31 | [-0.32, 0.91] |
| 16. Upper Division[ai] | 0.26 | 0.18 | [-0.09, 0.62] |
| 17. Low Enrollment[aj] | 0.01 | 0.22 | [-0.42, 0.44] |
| 18. Medium Enrollment[aj] | 0.14 | 0.16 | [-0.17, 0.46] |
| 19. Stand Alone[ak] | 0.16 | 0.28 | [-0.39, 0.71] |
| 20. Biology[ak] | -0.04 | 0.22 | [-0.46, 0.39] |
| 21. Hybrid[akl] | -0.09 | 0.18 | [-0.46, 0.27] |
| 22. Modality DNR[al] | -0.52 | 0.54 | [-1.58, 0.55] |

Notes: Estimated coefficients of variables for Δ intention were calculated using ordinary least squares (OLS) method considering that the intraclass correlation coefficient (ICC) was not significantly different from zero. Statistical estimates indicate the effects of predictors on the outcome for the focal group as compared to the reference group. [a]Effect coded such that the reference group = -0.5, and the focal group = 0.5. [b]Group-mean centered variable. [c]Grand-mean centered variable. [d]Reference group = continuing-generation college student. [e]Reference group = racial majority student. [f]Reference group = female. [g]Reference group = first-year student. [h]Reference group = combined mode (lab and lecture). [i]Reference group = lower division. [j]Reference group = high-enrollment course. [k]Reference group = integrated into microbiology course. [l]Reference group = in-person.

**Table S5. Longitudinal measurement invariance tests**

| *Model* | $\chi^2$ | *df* | *CFI* | *SRMR* | *RMSEA* | *90% CI* | *Model Comparison* | *Pass?* |
|---|---|---|---|---|---|---|---|---|
| | <u>Science Career Persistence Intentions</u> | | | | | | | |
| 1.1 Configural[A] | 533.976*** | 69 | .913 | .074 | .099 | [.092, .107] | -- | -- |
| 1.2 Configural[B] | 254.008*** | 59 | .964 | .055 | .070 | [.061, .079] | -- | -- |
| 1.3 Metric | 262.478*** | 65 | .963 | .057 | .067 | [.058, .075] | 1.2 vs. 1.3 | Y |
| 1.4 Scalar | 289.083*** | 71 | .959 | .058 | .067 | [.059, .075] | 1.3 vs. 1.4 | Y |
| | <u>Scientific Self-Efficacy</u> | | | | | | | |
| 2.1 Configural | 1.235 | 5 | 1.000 | .006 | .000 | [.000, .010] | | -- |
| 2.2 Metric | 4.660 | 7 | 1.000 | .016 | .000 | [.000, .036] | 2.1 vs. 2.2 | Y |
| 2.3 Scalar | 4.866 | 9 | 1.000 | .016 | .000 | [.000, .024] | 2.2 vs. 2.3 | Y |
| | <u>Scientific Identity</u> | | | | | | | |
| 3.1 Configural[A] | 85.857*** | 15 | .981 | .028 | .083 | [.066, .100] | | -- |
| 3.2 Configural[B] | 40.684** | 13 | .992 | .019 | .056 | [.037, .075] | | -- |
| 3.3 Metric | 45.399** | 16 | .992 | .025 | .052 | [.034, .070] | 3.2 vs. 3.3 | Y |
| 3.4 Scalar | 83.886*** | 19 | .982 | .030 | .070 | [.055, .086] | 3.3 vs. 3.4 | N |
| 3.5 Partial Scalar | 52.684*** | 18 | .990 | .027 | .053 | [.037, .070] | 3.3 vs. 3.5 | Y |
| | <u>Scientific Community Values</u> | | | | | | | |
| 4.1 Configural | 29.794* | 15 | .995 | .014 | .038 | [.017, .057] | | -- |

| | | | | | | | | |
|---|---|---|---|---|---|---|---|---|
| 4.2 Metric | 31.270* | 18 | .996 | .016 | .033 | [.011, .051] | 4.1 vs. 4.2 | Y |
| 4.3 Scalar | 33.082* | 21 | .996 | .016 | .029 | [.004, .047] | 4.2 vs. 4.3 | Y |

Note: Configural[A] = simple residual matrix; Configural[B] = complex residual matrix
*$p$ < 0.05, **$p$ < 0.01, ***$p$ <0.001.

**Table S6. Cross-sectional measurement invariance test between each group based on gender or HEC status**

| *Model* | $\chi^2$ | *df* | *CFI* | *SRMR* | *RMSEA* | *90% CI* | *Model Comparison* | *Pass?* |
|---|---|---|---|---|---|---|---|---|
| Gender Status T1 | | | | | | | | |
| 5.1 Configural | 545.899*** | 372 | .973 | .055 | .045 | [.037, .053] | -- | -- |
| 5.2 Metric | 579.618*** | 400 | .972 | .061 | .044 | [.036, .052] | 5.1 vs. 5.2 | Y |
| 5.3 Scalar | 632.468*** | 436 | .970 | .062 | .044 | [.037, .052] | 5.2 vs. 5.3 | Y |
| Gender Status T2 | | | | | | | | |
| 6.1 Configural | 722.087*** | 372 | .954 | .062 | .065 | [.057, .072] | -- | |
| 6.2 Metric | 757.789*** | 400 | .953 | .070 | .063 | [.056, .070] | 6.1 vs. 6.2 | Y |
| 6.3 Scalar | 810.193*** | 436 | .950 | .070 | .062 | [.055, .068] | 6.2 vs. 6.3 | Y |
| HEC Status T1 | | | | | | | | |
| 7.1 Configural | 402.358*** | 248 | .976 | .052 | .043 | [.035, .050] | -- | |
| 7.2 Metric | 425.218*** | 262 | .975 | .059 | .043 | [.035, .050] | 7.1 vs. 7.2 | Y |
| 7.3 Scalar | 465.867 | 280 | .971 | .057 | .044 | [.037, .057] | 7.2 vs. 7.3 | Y |
| HEC Status T2 | | | | | | | | |
| 8.1 Configural | 550.733*** | 248 | .959 | .053 | .060 | [.053, .067] | -- | |
| 8.2 Metric | 564.883*** | 262 | .959 | .056 | .058 | [.052, .065] | 8.1 vs. 8.2 | Y |
| 8.3 Scalar | 596.605*** | 280 | .958 | .055 | .058 | [.051, .064] | 8.2 vs. 8.3 | Y |

Note: Gender status comparisons were conducted across 3-groups: Female, Male, and Other/Did not report. HEC status comparisons were conducted across 2-groups: historically excluded racial/ethnic communities in STEM and majority groups in STEM.
*$p < 0.05$, **$p < 0.01$, ***$p < 0.001$.
